# Quantitative interactome mapping of skeletal muscle insulin resistance

**DOI:** 10.64898/2026.01.05.697787

**Authors:** Yaan-Kit Ng, Ronnie Blazev, Julian P. H. Wong, Nichollas E. Scott, Jeffrey Molendijk, Benjamin L. Parker

## Abstract

Protein-protein interactions (PPIs) are dynamic and critical to adaptive homeostasis. While there have been massive efforts to catalogue proteome-wide PPIs, global quantification of changes remains a challenge. Here, we integrate dynamic protein correlation profiling - mass spectrometry (PCP-MS) and quantitative cross linking-mass spectrometry (qXL-MS) using multiplexed stable isotope labelling to characterise global PPI remodelling following the development of chronic skeletal muscle insulin resistance (IR) with or without acute insulin stimulation. We quantify >7,000 unique PPIs amongst 5,346 proteins and show changes in the interactome network dominate the proteome response. Our data show the dysregulation of protein processing in the endoplasmic/sarcoplasmic reticulum involving changes in PPIs with protein chaperones and disulfide isomerases is a major hallmark of skeletal muscle IR. Mechanistically, we show the dysregulation of PPIs with Protein-Disulfide Isomerase 6 (PDIA6) regulates cysteine oxidation and insulin sensitivity. Taken together, we show *in vivo* quantitative interactome mapping is a powerful approach to understand disease mechanisms and provide new insights into protein network re-organisations with IR.

## Introduction

Protein-protein interactions (PPIs) are essential for protein function that fine tune physiological homeostasis. It is therefore not surprising that characterizing proteome-wide changes in PPIs has important implications to understand health and disease (Greenblatt et al., 2024, Liu et al., 2026). The expansion of mass spectrometry (MS)-based proteomics has seen the development of several methods for large-scale discovery and quantification of PPIs. Affinity purification-mass spectrometry (AP-MS) incorporating exogenously tagged proteins or immunoprecipitation (IP) of endogenous complexes have identified thousands of interactions across a variety of cell types (Huttlin et al., 2017, Oughtred et al., 2021, Pankow et al., 2016, Lambert et al., 2013). However, limitations include the requirement to target individual proteins, and weak or transient proteins may be lost during sample preparation. Furthermore, enrichment of entire protein complexes and high-throughput analysis typically cannot distinguish between direct or indirect interactions. Protein correlation profiling-mass spectrometry (PCP-MS) incorporating the biophysical separation of protein complexes followed by identification of co-eluting proteins is an orthogonal approach to identify proteome-wide PPIs (Kristensen et al., 2012, Bludau et al., 2020, Rosenberger et al., 2020, Skinnider et al., 2021, Schulte et al., 2023, Larance et al., 2016, Palenikova et al., 2023, Havugimana et al., 2022). These approaches avoid the need to tag and/or enrich specific protein complexes making them more suitable for *in vivo* analysis however, they generally require lysis in native buffers which may not allow efficient extraction of e.g. membrane proteins, and similar to AP-MS, they cannot validate direct interactions (Babu et al., 2012). Despite this, the use of PCP-MS combined with stable isotope labelling or label-free quantification is a powerful approach to characterise dynamic changes in PPIs (Heusel et al., 2019, Kristensen et al., 2012).

Cross linking-mass spectrometry (XL-MS) has grown to be a popular method that overcomes several limitations mentioned above by introducing a cross linker to covalently bind interacting proteins. A variety of cross linkers have been developed with defined distance constraints which can be applied to live cells and react with proteins within direct proximity. Protein complexes can be extracted with harsh denaturing buffers and following protease digestion, XL-peptides can be identified by MS to infer native direct protein-protein interactions (Piersimoni et al., 2022). However, the identification of XL-peptides has additional considerations as they are typically less abundant than non-XL-peptides in a complex reaction mixture and require additional enrichment/separation techniques for proteome-wide analysis. Furthermore, several acquisition and data analysis considerations are unique to XL-peptide characterisation such as the requirement to determine the sequences of multiple peptides within individual MS/MS or MS^n^ scans (Leitner et al., 2020, Yilmaz et al., 2018, Iacobucci et al., 2019). Several groups have performed cross linking of live cells to capture PPI landscapes with the largest datasets identifying over 6,000 unique protein interactions (Wheat et al., 2021, Brauer et al., 2025, Bartolec et al., 2023, Zhang et al., 2025, Gao et al., 2022, Burke et al., 2023). Furthermore, recent advances have enabled quantification of XL peptides to gain further biological insight beyond simple identification. Although label-free approaches are feasible, cross-linking studies commonly use stable isotope tagging—either built into the cross-linker itself (Chen et al., 2016, Muller et al., 2001, Chavez et al., 2020) or labelling the peptide (Yu et al., 2016, Ruwolt et al., 2022). However, most quantitative applications have focused on conformational analysis (Keller et al., 2025, Yu et al., 2016) based on the idea that XL-peptides spanning flexible regions will change in abundance following changes in protein structure. Another forefront that remains limited is the application of XL-MS on whole tissue due to the increased complexity of tissues containing multiple cell types and extracellular matrix, and the limited ability of cross linkers to permeate the entire sample to effectively capture the native interactome (Chavez et al., 2018). To date, only a limited number of studies have performed qXL-MS of tissue samples where Caudal et al. (2022) performed *ex vivo* crosslinking of healthy or failing mouse hearts diced into ∼1mm^3^ pieces followed by mitochondrial isolation to quantify 3,792 XL-peptides from ∼500 PPIs. Furthermore, Bakhtina et al. (2023) crosslinked isolated mitochondria from gastrocnemius muscle to identify aging associated changes in PPIs (Caudal et al., 2022, Bakhtina et al., 2023). While this reflects the challenging nature of whole tissue crosslinking, the benefit of this approach when also coupled with qXL-MS has exciting potential to characterise the remodelling of PPIs during disease progression.

In this study, we established a whole tissue qXL-MS workflow to investigate proteome-wide changes in PPIs in mouse skeletal muscle insulin resistance (IR). These data were integrated with a quantitative analysis of IR in differentiated C2C12 myotubes maintained in cell culture using both PCP-MS and qXL-MS. We also included the quantification of dynamic changes in PPIs when control or IR myotubes were treated with or without acute 30 minutes of insulin stimulation and show that PCP-MS outperforms qXL-MS to capture dynamic changes in acute remodelling. The workflows incorporated the quantification of total protein levels to pinpoint changes in proteome abundance vs PPI remodelling during the development of IR. Our integrated analysis has identified hundreds of changes in PPI’s following the induction of IR and we further validate PDIA6, a chaperone and cysteine redox modification isomerase that is a central hub in the interaction network. *In vivo* over-expression of PDIA6 modulated insulin-stimulated glucose uptake, and redox proteomics quantified changes in precise cysteine modification sites on interacting proteins.

## Results

### Probing acute insulin signalling protein complex dynamics in insulin resistant C2C12 myotubes with PCP-MS

To study dynamic remodelling of protein complexes, C2C12 myotubes were treated with 0.5 mM palmitic acid for 24 h to induce insulin resistance (IR) and then stimulated with or without acute 100 nM insulin for 30 min followed by PCP-MS (**Fig. 1A**) (n=3 each). Western blotting confirmed activation of the insulin signalling pathway following acute stimulation including phosphorylation of S473 on Akt and its direct substrate S580 on AS160, and this was attenuated in the IR group (**Fig. 1B-C**). Samples were lysed in non-denaturing buffer, separated by BN-PAGE, and each lane cut into 30 x 1.5mm fractions. Each fraction underwent in-gel digestion and was analysed with liquid chromatography – tandem mass spectrometry (LC-MS/MS) using data-independent acquisition (DIA) resulting in the analysis of 360 individual runs. Principal component analysis (PCA) revealed separation of IR in PC1 followed acute insulin stimulation in PC2 (**Fig. 1D**). The PCP-MS data were analysed by CCProfiler which allows the identification of protein complex re-organisation in quantitative co-fractionation experiments (Bludau et al., 2023) resulting in a total of 7,672 proteins identified with 4,680 passing filtering (see Methods) (**Supplementary Table S1**). We first performed a complex-centric analysis by mapping proteins to the CORUM, STRING and Hu.MAP3 databases to retrieve known PPIs (Fischer et al., 2024). This identified 166 protein complex features defined as having two or more subunits of a defined complex sharing a coelution peak (see Methods for filtering) (**Supplementary Table S2**). **Fig. 1E** shows examples of protein complexes displaying high co-elution correlations including the CCT Complex, Proteasome, Prefoldin Complex, WASH Complex and AP-4 Complex. Next, we investigated the regulation of these known protein complexes in response to IR with or without acute insulin stimulation. This included two analyses: i) a global analysis by taking the area under the curve across the entire elution profile to calculate changes in the overall abundance which is particularly important for the analysis of chronic IR over several hours which may induce changes in protein expression levels, and ii) local analysis by taking the area under the curve across a local window to investigate how subunits of protein complexes are assembled or disassembled with a particular focus on how these may change following acute insulin stimulation in healthy and disease settings. Out of the 166 protein complexes, 17 local complex features changed with insulin stimulation in the healthy cells with only 5 changing in IR myotubes suggesting impaired re-organisation of protein complexes with onset of IR. Interestingly, there were 47 local features that changed between basal states of healthy and IR myotubes, and 41 features that changed between stimulated states of healthy and IR myotubes suggesting most changes occur with onset of IR as opposed to acute insulin stimulation.

**Figure 1.**
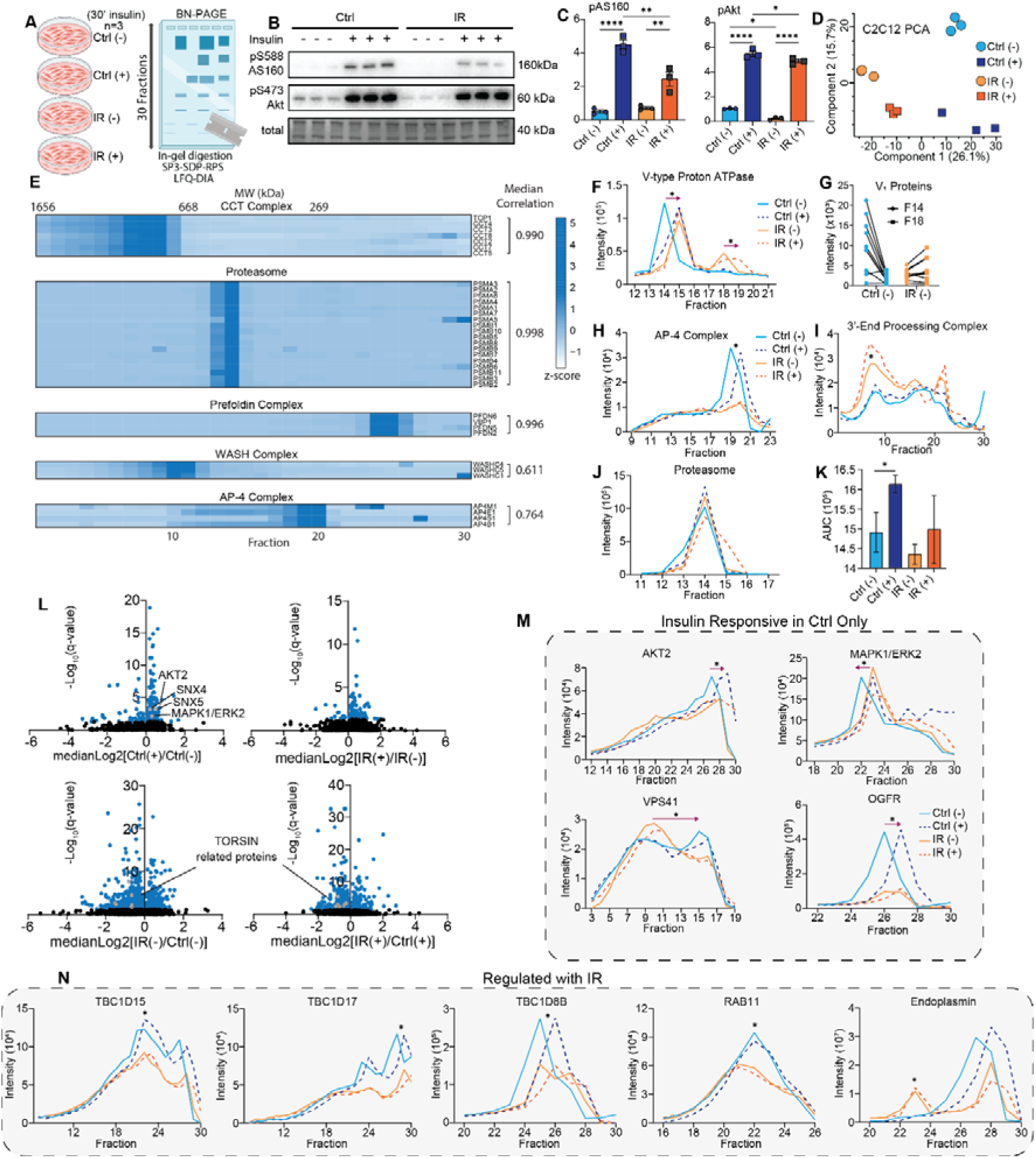
Protein Correlation Profiling (PCP)-MS analysis of healthy and insulin resistant C2C12 myotubes. (**A**) Experimental workflow detailing PCP-MS of healthy and insulin resistant C2C12 myotubes (n=3) that were either basal or stimulated with insulin. (**B**) Western blot, and (**C**) densitometry quantification of Akt and AS160 phosphorylation. (**D**) Principal component analysis. (**E**) Heatmap representing the intensity of protein complexes and their subunits across the fractions. (**F**) Elution profiles of V-type Proton ATPase, and (**G**) quantification of V1 subunits. Elution profiles of (**H**) AP-4 Complex, (**I**) 3’-End Processing Complex, and (**J**) Proteasome with quantification shown in (**K**). (**L**) Volcano plot visualizing all identified local protein features and their regulation in response to both insulin stimulation and development of IR. (**M**) Elution profiles of protein features that are regulated with insulin stimulation in the control C2C12 myotubes only. (**N**) Elution profiles of protein features regulated with onset of IR. *p<0.05, **p<0.01, ****p<0.0001, unpaired t-test. Bars depict mean values and error bars represent SEM.

This complex centric analysis identified several features that were regulated with either insulin stimulation or with induction of IR. For example, the V-type proton ATPase complex assembly was identified as a local feature residing at approximately 720kDa, which aligns with its expected molecular weight (MW) (Wang et al., 2020) (**Fig. 1F**). In response to acute stimulation there was a shift towards a lower molecular weight suggesting partial disassembly, while induction of IR reduced the complex into two distinct peaks assembled at ∼700 kDa (Fraction 14) and ∼550 kDa (Fraction 18). V-ATPase is known to govern the trafficking of the fatty-acid transporter CD36, which is typically sequestered in endosomes that require luminal acidification by V-ATPase. Excess lipid exposure has been shown to cause the V-ATPase to disassemble into its V_1_ and V_0_ subcomplexes, raising endosomal pH and triggering CD36 translocation to the plasma membrane (Liu et al., 2017). This increases fatty acid uptake while activating pro-inflammatory Toll-like receptor-4 (TLR4) signalling, promoting lipid overload, inflammation, and insulin resistance (Wang et al., 2024). This disassembly is reflected in our data as we see a peak at ∼550 kDa consisting of only the V_1_ subcomplex proteins (**Fig. 1G**). Induction of insulin resistance in C2C12 myotubes also drove disassembly of the adaptor protein complex-4 (AP-4) (**Fig. 1H**). AP-4 is a heterotetrameric vesicle coat complex localized to the trans-golgi network involved in sorting cargo proteins to endosomes and other destinations. AP-4 has been shown to regulate autophagy via trafficking of the autophagy protein ATG9A (Davies et al., 2018) where impaired autophagy in skeletal muscle is known to influence insulin sensitivity (Frendo-Cumbo et al., 2019). Hence, these data provide important mechanistic clues about the regulation of autophagy in IR skeletal muscle potentially via changes in AP-4 PPIs. In contrast to changes in assembly/disassembly, the 3’-end processing complex showed increased global abundance with insulin resistance (**Fig. 1I**). The 3′-end processing complex regulates the cleavage and polyadenylation of the 3′ end of pre-mRNAs where recent evidence suggests that alterations in mRNA 3′-end processing, particularly alternative polyadenylation, can reprogram gene expression in metabolic tissues and contribute to insulin resistance (Li et al., 2015). In response to acute insulin stimulation, there was also increased complexing of the 20S proteasome independent of global abundance, a response that was not observed under IR conditions (**Fig. 1J-K**). This is interesting as acute insulin stimulation is typically associated with a reduction in the expression of protein degradation genes (Kaiser et al., 2022). However, prolonged mTORC1 activation has been shown to trigger a coordinated increase in proteasome production in mouse skeletal muscles as well as mouse embryonic fibroblasts (MEFs) (Kaiser et al., 2022, Zhang et al., 2014). It is important to note however, that these studies have typically investigated expression/abundance and do not specifically characterize complex dynamics.

We next focused our analysis on a protein centric approach. This enabled the identification of protein interaction changes independent of pre-defined complexes. This analysis was similar to the complex-centric method but used peptide intensities to quantify proteins instead of subunit intensities for complexes. This revealed 3,698 protein features, with 316 and 141 local features significantly regulated by insulin in healthy and IR myotubes, respectively (**Supplementary Table S3**). Additionally, IR regulated 882 local features in the basal state and 699 in the stimulated state (**Fig. 1L**). Initially, we interrogated proteins significantly shifting in response to insulin stimulation in the control myotubes (**Fig. 1M**). Among these were canonical insulin signalling proteins including AKT2 and ERK2 which saw changes in local protein features in healthy controls only. Vesicle trafficking, endosomal recycling and cellular sorting associated proteins also had a similar response profile. For example, an identified local protein feature of VPS41 around ∼400 kDa (Fraction 15/16) had reduced abundance in response to insulin stimulation in the healthy myotubes only (**Fig. 1M**). There was also a clear shift in complex formation between healthy and IR states with a greater peak observed around fraction 9 in the IR state. VPS41 is a subunit of the HOPS (homotypic fusion and vacuole protein sorting) tethering complex, classically required for fusion of late endosomes with lysosomes. The molecular weight of HOPS is approximately 400kDa and aligns with the feature identified in fraction 15/16. This complex is also known to govern endo-lysosomal trafficking and autophagic flux (Jiang et al., 2024, Liu et al., 2023). There was also an insulin stimulated shift towards a heavier molecular weight for a protein feature identified in OGFR. This response is not only absent in IR, but OGFR is also significantly reduced in global abundance. OGFR has emerged as a regulator of adipose tissue function in obesity where adipocyte-specific OGFR ablation results in reduced energy expenditure and exacerbates the development of insulin resistance when on a high-fat diet (Zhang et al., 2023).

The final CCProfiler analysis focused on protein features that significantly change with the onset of IR (**Fig. 1N**). This includes an array of TBC1 domain-containing proteins (TBC1D15, TBC1D17, TBC1D8B) that are consistently reduced with IR. Small GTPases of the Rab family orchestrate vesicle trafficking, while TBC1Ds act as Rab GTPase-activating proteins (GAPs) to modulate these processes (Cartee, 2015). TBC1D15 is a Rab-GAP originally identified to act on the late endosomal/lysosomal Rab7, and emerging evidence links it to GLUT4 vesicle trafficking where Wu et al. (2019) reported that TBC1D15 knockout cells show a significant reduction in glucose uptake (measured by 2-NBDG uptake) alongside a decrease in total GLUT4 protein content (Wu et al., 2019). TBC1D17 has recently been identified as a RAB5-specific GAP that connects AMPK to glucose transport where RAB5 governs early endosome fusion and has also been shown to participate in GLUT4 vesicle sorting and insulin signalling (Rao et al., 2021). TBC1D8B is a less-studied TBC1D protein that has gained attention due to its link with RAB11 which also shows a similar reduction in local feature abundance with IR. It has been shown in podocytes that TBC1D8B binds preferentially to the active form of RAB11 and likely functions to inhibit RAB11 signalling (Kampf et al., 2019). RAB11 is a master regulator of slow recycling endosomes, which return internalized proteins to the plasma membrane and redistribute intracellular membranes (Zulkefli et al., 2019). Furthermore, Uhlig et al. showed that in cardiac muscle cells and skeletal muscle myotubes, a dominant-negative RAB11 mutant reduced insulin-stimulated GLUT4 translocation and glucose uptake by about 50% (Uhlig et al., 2005). Many protein features regulated with IR also belonged to several ER associated proteins. This included TORSIN related proteins (TOR1AIP1, TOR1AIP2, TOR1B, TOR3A) which all had significantly reduced local abundance with IR (**Fig. 1L**). TORSINs are the only AAA+ ATPases that reside in the nuclear envelope and ER (Rampello et al., 2020). TORSINA is the best-characterized isoform, activated by cofactors LAP1 (TOR1AIP1) and LULL1 (TOR1AIP2), which are membrane-bound regulators in the inner nuclear membrane and ER, respectively (Rose et al., 2015). While their specific functions remain elusive, recent mouse studies showed that loss of TORSINA in the liver causes hepatic steatosis and ER lipid droplet accumulation where combined deletion of LAP1 and LULL1 recapitulates the TORSINA knockout phenotype, emphasizing their essential co-regulatory function (Hernandez-Ono et al., 2024). Furthermore, the elution profile of Endoplasmin reveals both an insulin stimulated shift in a feature with an apex at Fraction 28, as well as an appearance of a protein assembly in the IR state only at fraction 23. Endoplasmin (also known as GRP94 or HSP90B1) is a glucose-regulated ER chaperone essential for folding secreted and membrane proteins and is involved in the unfolded protein response (Malhotra & Kaufman, 2007). Additionally, macrophage specific KO of GRP94 was found to improve glucose tolerance and insulin sensitivity while lowering macrophage numbers to reduce chronic inflammation (Song et al., 2020). Our data support widespread changes in PPIs at the ER with the induction of IR. Taken together, we present dramatic remodelling of protein complexes during palmitate-induced IR and show how acute insulin stimulation can fine tune PPIs.

### Validating the regulation of PPIs with insulin resistance in C2C12 myotubes with quantitative XL-MS

We next sought to validate changes in direct physical PPIs using qXL-MS. Here, we used the well-established cell permeable and enrichable cross-linker t-butyl-PhoX (tPhoX) (Jiang et al., 2022). C2C12 myotubes were made IR as above with 0.5 mM palmitic acid for 24 h along with non-treated controls and then each group stimulated with or without 100 nM insulin for 30 min (n=4 each) (**Fig. 2a**). Cells were then cross-linked with 2 mM of tPhoX for a further 30 min in the presence of insulin and the reaction quenched as previously described (Jiang et al., 2022). To increase the depth of analysis, each sample was subjected to a crude biochemical-based separation into four fractions (Wong et al., 2025) followed by digestion with trypsin/LysC. The 16 samples from each fraction were labelled with 16-plex Tandem Mass Tags (TMT) and pooled, followed by the depletion of phosphopeptides and enrichment of XL-peptides using immobilised metal-ion affinity chromatography (IMAC). Each of the four biochemical fractions underwent peptide-level size-exclusion chromatography (SEC) to partially fractionate larger XL-peptides from mono-/loop-linked peptides in 2-3 fractions which were further separated by high pH reversed-phase chromatography into 12 fractions and analysed by LC-MS/MS using data-dependent acquisition (DDA) resulting in the analysis of 132 individual runs. Data were analysed with pLink2 and searched against a skeletal muscle-specific mouse proteome database to limit the search space with filtering to 1% FDR followed by reporter-ion quantification and statistical analysis (see Methods). Prior to this analysis, we first optimised the collision energies for the analysis TMT-labelled XL-peptides using tPhoX-reacted BSA and found that a stepped collision energy of 35 ± 8 produced the greatest number of cross-linked spectral matches (CSMs) and richer fragment-ion information with the largest average MS/MS identification scores (**Fig. 2B**). A total of 88,829 unique peptides were identified of which 61,691 were mono-linked, 10,645 were loop-linked, 11,295 were XL and 5,198 were regular non-PhoX modified peptides (**Fig. 2C**) (**Supplementary Table S4**). The XL-peptides were made up of 6,351 intra-protein and 4,944 inter-protein crosslinks corresponding to 3,757 unique protein pairs. Principal component analysis (PCA) showed distinct separation between control and IR cells however insulin stimulation did not drive significant separation in the crosslinked proteome **(Fig. 2D**). These data suggest that unlike the above PCP-MS, qXL-MS was not able capture acute dynamic changes in PPIs under these experimental conditions. Hence, for the remainder of the analysis we grouped the insulin-stimulated and non-simulated samples and focused on control vs IR comparison (n=8 each). A change in abundance of a XL peptide could be due to changes in PPIs or be driven by global changes in the abundance of one or both interacting proteins. Hence, we performed a total proteomic analysis using an aliquot of peptides that did not bind to IMAC. This analysis identified 7,850 proteins where 2,989 were regulated with IR (**Supplementary Table S5**). Next, each XL peptide was mapped to the corresponding abundance of each interacting protein to investigate changes in XL-peptide abundance vs protein abundance. This was visualized with two plots where the x-axis shows the Log_2_ ratio (IR/control) of the XL peptide while the y-axis shows the ratio of either Protein 1 or Protein 2 in the complex (**Fig. 2E-F**). A total of 6,704 out of 11,295 XL peptides were significantly regulated in IR C2C12 myotubes where 5,214 XL peptides also changed without a shift in abundance of either interacting partner. A KEGG pathway analysis was performed using the proteins containing regulated XL peptides revealing an enrichment of protein interactions associated with vesicular transport, ribosome, fatty acid metabolism, protein processing in endoplasmic reticulum (ER), proteasome and proteolytic activity, thermogenesis, endocytosis and calcium signalling (**Fig. 2G**).

**Figure 2.**
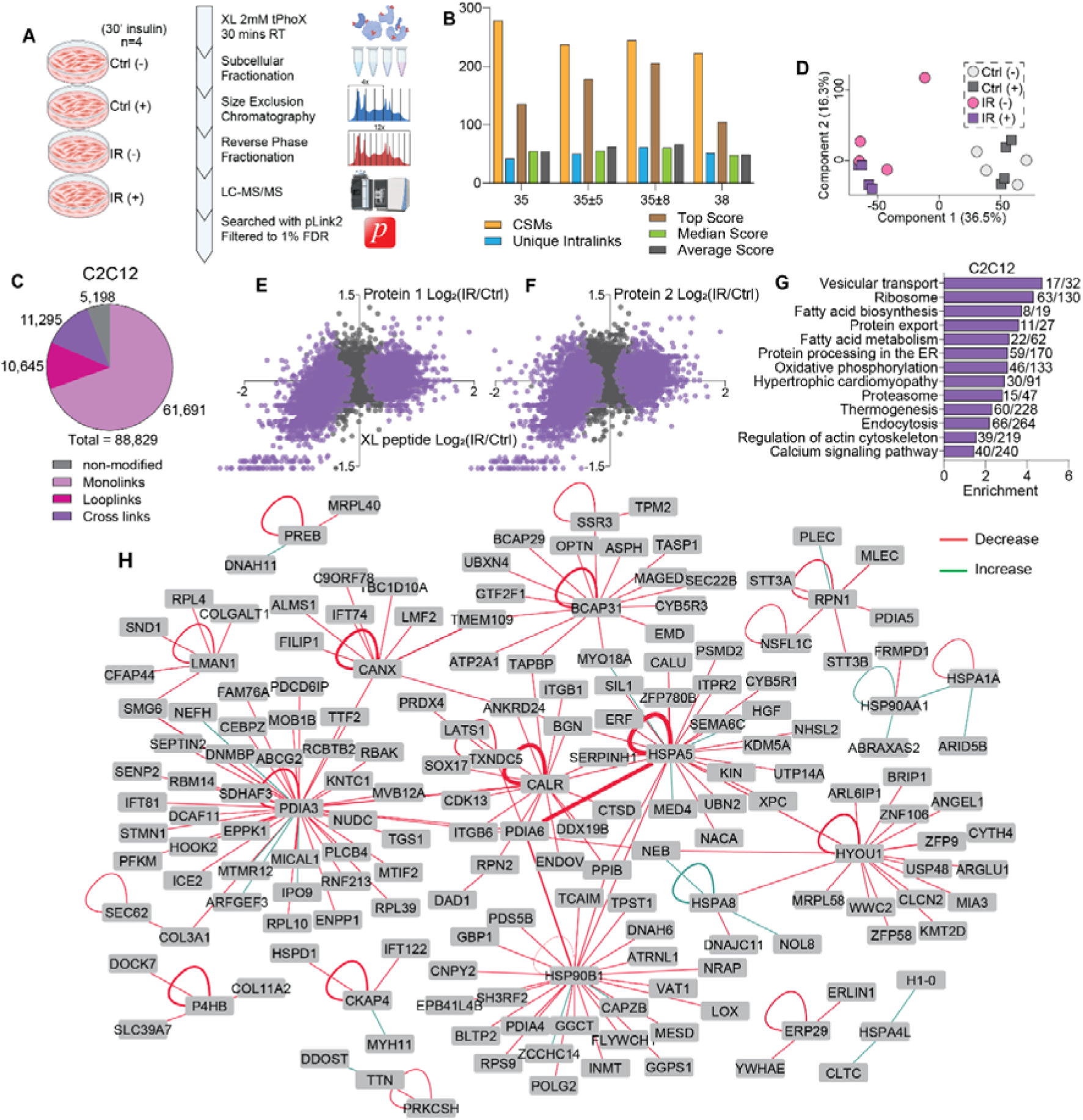
Cross linking (XL)-MS analysis of healthy and insulin resistant C2C12 myotubes. (**A**) Experimental workflow detailing XL-MS of healthy and insulin resistant C2C12 myotubes (n=4) that were either basal or stimulated with insulin. (**B**) Crosslink spectral matches across several normalised collisional energies. (**C**) Comparison of regular, mono linked, loop linked and cross-linked peptides identified in this dataset. (**D**) Principal component analysis of each biological group. (**E-F**) A scatterplot comparing the fold change of each cross linked (XL) peptide to the fold change of each interacting protein between IR and control samples. (**G**) KEGG pathways enriched in significantly regulated XL peptides. (**H**) XL-peptide interactome for ER protein processing where edge thickness shows cross-link count and edge colour marks change in IR myotubes vs control (green increase, red decrease).

Protein processing in the ER was particularly prominent because in addition to the KEGG enrichment, the pathway had very high representation of the total number of XL-peptides significantly regulated and also had a high degree of total proteome regulation. These data are consistent with the protein-centric PCP-MS analysis which also showed changes in proteins involved in proteostasis at the ER. **Figure 2H** depicts an interactome map of XL proteins involved in protein processing in the ER where the thickness of the edges represents the number of cross links between each node while the colour indicates either an increased (green) or decreased (red) fold change with IR. There were several cross links identified between a family of protein disulfide isomerases (PDIA1, PDIA3, PDIA4 and PDIA6). HYOU1, a molecular chaperone upregulated during ER stress, presented with various cross links which was interesting given its overexpression has been shown to improve insulin sensitivity and glucose handling (Nakatani et al., 2005). Several cross links were also identified with calreticulin which is a known ER-localized protein able to bind Ca2+ and misfolded proteins (Jeffery et al., 2011). Similarly, there were several inter-protein crosslinks identified with HSPA5 (BiP), a master regulator of the UPR, that were reduced with IR (Pobre et al., 2019). Taken together, our proteome-wide qXL-MS of C2C12 myotubes serves as valuable tool to validate novel PPIs and has revealed a vast interaction landscape remodelled with IR with major potential defects in protein folding and chaperone binding.

### *Ex vivo q*uantitative XL-MS of mouse skeletal muscle insulin resistance

While our PCP-MS and qXL-MS of IR in C2C12 myotubes contains hundreds of validated PPIs, it is limited to only an *in vitro* setting. Hence, the major goal of the current study was to establish a whole tissue skeletal muscle qXL-MS workflow to study remodelling of PPIs with IR. Mice were feed a standard chow diet or a high fat diet (HFD) for 12 weeks which induced skeletal muscle insulin resistance as assessed by attenuated *ex vivo* insulin-stimulated ^3^H-2-deoxyglucose uptake into *Soleus* muscle and reduced phosphorylation of pS473 on Akt and pS588 on AS160 (**Fig. 3A-C**). The *Extensor digitorum longus* (EDL) skeletal muscles were isolated in parallel from the identical mice and subjected to qXL-MS analysis. We chose this small fast-twitch muscle because it weighs ∼8-10 mg providing sufficient protein yield (∼1 mg) and it is commonly used in *ex vivo* myograph preparations due to adequate oxygen diffusion in O_2_/CO_2_-carbogen gassed Krebs-Henseleit bicarbonate-based buffer (Barclay, 2005). We first optimised *ex vivo* tPhoX cross-linking of EDL and found that a maximum concentration of 10 mM tPhoX for 2h in carbogen bubbled modified Krebs-Henseleit Buffer at 30°C produced the greatest number of XL peptide identifications (**Fig. 3D**). This timeframe has been extensively used in metabolic and/or contractile functional assessments (Hansen et al., 1994, Ryder et al., 2000, Jakobsgaard et al., 2021). Analysis by SDS-PAGE revealed subtle shifts to heavy molecular weights suggesting samples were not over cross-linked and are comparable to previously published cell-based cross-linking protocols (Jiang et al., 2022, Liu et al., 2015, Wheat et al., 2021, Kaake et al., 2014) (**Fig. 3E**). For XL-MS analysis of IR in skeletal muscle, chow and HFD EDL muscles (n=8 each) were incubated with 10 mM tPhoX for 2h. Unlike our analysis of C2C12 myotubes, we did not include an analysis of acute insulin stimulation. Each of the 16 muscles underwent the identical workflow as performed on C2C12 myotubes with biochemical subcellular fractionation into four fractions followed by TMT labelling, IMAC enrichment, SEC fractionation and high pH fractionation followed by LC-MS/MS and analysis by pLink2 against the skeletal muscle-specific mouse proteome database using as described above. We identified a total of 72,120 peptides of which 52,515 were mono-linked, 7,186 were loop-linked, 8,327 were XL and 4,092 were regular non-modified peptides (**Fig. 3F**) (**Supplementary Table S6)**. The XL-peptides were made up of 2,777 intra-protein and 5,550 inter-protein crosslinks corresponding to 3,541 unique protein pairs. PCA of the XL peptides revealed separation between the chow and HFD fed mice (**Fig. 3G**). Similar to the C2C12 analysis, a change in the abundance of a XL peptide may be driven by either a remodelling of PPIs or by changes in the abundance of the interacting proteins themselves. Therefore, we deconvoluted this by performing a total proteomic analysis using an aliquot of TMT-labelled peptides prior to IMAC enrichment (**Supplementary Table S7**). All 8,327 unique XL peptides identified had corresponding total protein quantification available allowing us to plot the XL peptide verses total levels of the interacting proteins (“Protein 1” and “Protein 2”) fold-changes (**Fig. 3H-I**). A total of 1,101 XL peptides were significantly regulated in HFD diet mice where a smaller proportion of XL peptides (766) significantly changed without a change in expression of either interacting partner. Focusing on inter-protein crosslinks that; i) have the greatest significant change and ii) are exclusive to the *EDL* dataset, this identified several novel interactions between proteins implicated in skeletal muscle health and disease (RALGAPA2, ALDOA, HRC, CKM, APIA1/2, ENO3, LDLRAP1, LARS1) (**Fig. 3J**). For example, EXOC3 is part of the exocyst complex which is required for insulin-stimulated GLUT4 trafficking (Fujimoto et al., 2019) however it has not been previously shown to interact with SERPINA3K which functions as a protective signalling factor against ischemia-induced cardiac injury (Wang et al., 2025). A novel interaction between ATP2A1 (SERCA1) and MAGEL2 was also significantly decreased with IR. Both proteins are localised to the SR/ER with the former regulating calcium handling critical for contractile function, and the latter regulating endosomal transport. Interestingly *MAGEL2* is in the Prader-Willi Syndrome imprinting locus and is associated with obesity and reduced muscle mass, and mice lacking *Magel2* have increased adiposity and reduced muscle function (Kamaludin et al., 2016).

**Figure 3.**
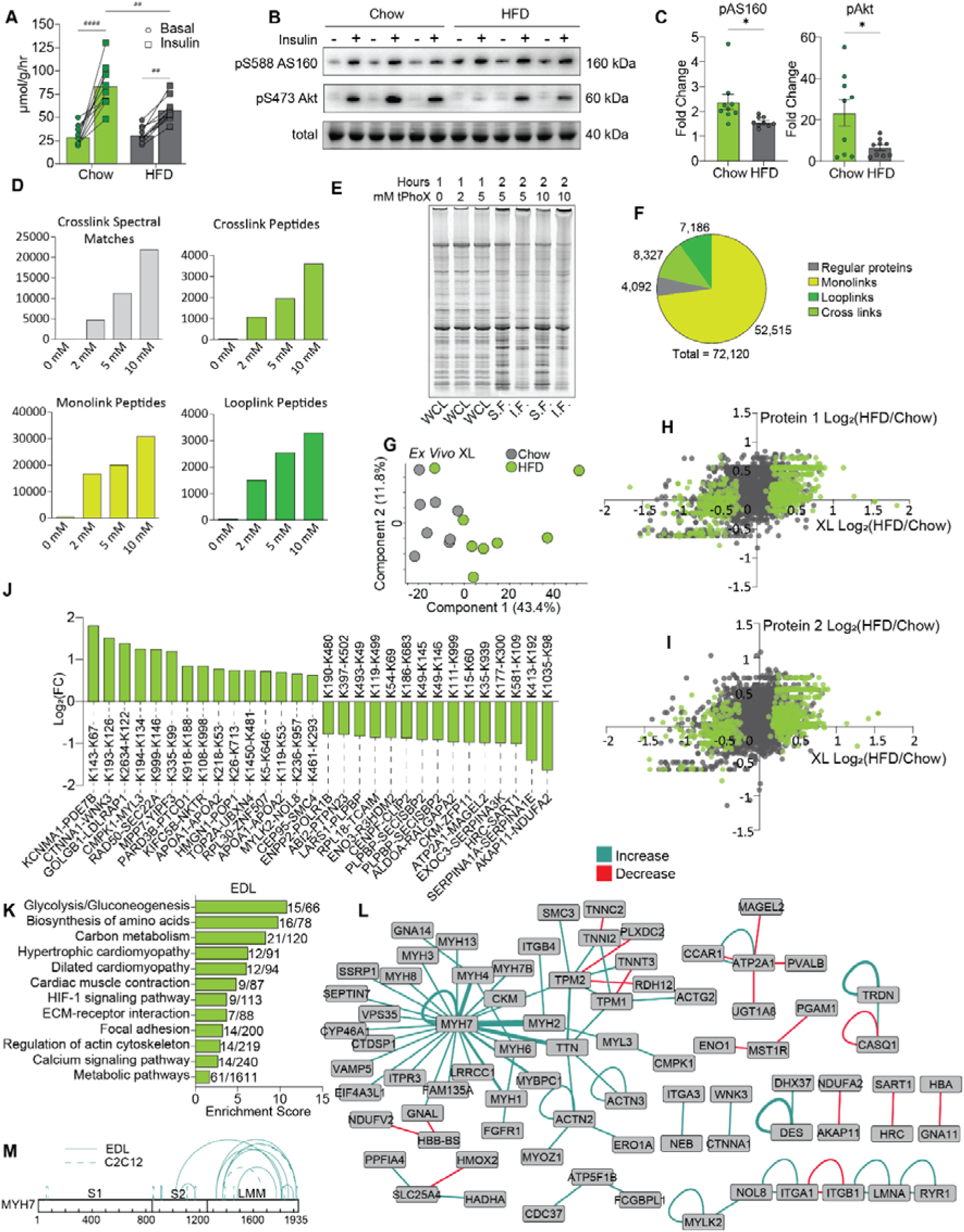
Cross linking (XL)-Mass Spectrometry analysis of Extensor Digitorum Longus of high fat diet (HFD) vs chow fed mice. (**A**) *Ex vivo* 90nM insulin stimulated 2-Deoxy-D-Glucose uptake in soleus muscles from chow and HFD fed mice. (**B**) Western blot analysis of chow and HFD soleus muscles investigating AS160 and Akt phosphorylation in response to 90nM insulin stimulation (n=9). (**C**) Quantification of soleus western blots. (**D**) Cross link spectral matches in muscle samples incubated across several crosslinker concentrations. (**E**) Total protein stain visualizing degree of crosslinking in muscles subject to various crosslinking conditions with and without crude fractionation. (**F**) Comparison of regular, mono linked, loop linked and cross-linked peptides identified in this dataset. (**G**) Principal component analysis of each biological group. (**H-I**) A scatterplot comparing the fold change of each cross linked (XL) peptide to the fold change of each interacting protein between HFD and chow samples. (**J**) Inter-protein XL peptides with the greatest fold change identified only in *EDL* (**K**) KEGG pathways enriched in significantly regulated XL peptides. (**L**) XL-peptide interactome for contractile and Ca^2+^ handling proteins where edge thickness shows cross-link count and edge colour marks change in IR myotubes vs control (green increase, red decrease). (M) Schematic of intra-protein XLs identified on MYH7. *p<0.05, **p<0.01, unpaired t-test. ##p<0.01, ####p<0.0001, paired t-test. Bars depict mean values and error bars represent SEM.

Performing KEGG pathway analysis on all significantly regulated XL peptides revealed increased representation of interactions associated with glycolysis/gluconeogenesis, carbon metabolism, cardiomyopathy, HIF-1 signalling, contractile function and calcium signalling, with the latter two partially interesting given their importance in muscle function and metabolism (**Fig. 3K)**. This included differentially regulated crosslinks identified between key calcium handling proteins including TRDN, CASQ1, ATP2A1, RYR1 and PVALB and several contractile apparatus proteins (MYH2, MYH4, MYL3, MYLK2, TPM1/2) (**Fig. 3L**). Interestingly, many of these highly regulated crosslinks are not associated with changes in protein abundance. Therefore, these changes in PPIs may underly the mechanisms of defective calcium handling observed in obese/diabetic models (Eshima et al., 2020, Bruton et al., 2002, Jain et al., 2014).

A striking observation was the large number of intralinks identified on Myosin 7 (MYH7) that were regulated with IR. This is interesting because the conformational changes in the relaxation state of myosin have been previously implicated the pathogenesis of cardiomyopathies as well as defective metabolism in muscle (Schmid & Toepfer, 2021, Carrington et al., 2023). Specifically, mutations and abnormal glycation events in the C-terminal light meromyosin (LMM) region of MYH7 are associated with changes in the relaxation state (Carrington et al., 2023, Lewis et al., 2025). In fact, a significantly greater proportion of MYH7 from type 2 diabetic (T2D) patients assume an energy-saving super relaxed state (SRX) instead of the energy inefficient disordered-relaxed state (DRX) which may contribute to metabolic imbalance. Integration with the C2C12 XL-MS data revealed that cross links regulated across both datasets concentrated on the LMM region of MYH7 (**Fig. 3M**). Hence, changes in the abundance of intralink peptides across the LMM region following the induction of IR may represent confirmational changes in myosin relaxation.

### Interactome integration

We first investigated the validity of our qXL-MS data by mapping XL peptides identified in C2C12 and/or EDL skeletal muscles onto experimentally resolved protein complex structures from the Protein Data Base (PDB). Here, we focused on only mouse structures and included only complexes with 3 or more full length subunits. This filtered our analysis to the Immunoproteasome, Mitochondrial Complex I, Actin Related Protein 2/3 Complex (ARP2/3 Complex), and the Actomyosin Complex captured in the rigor state (**Fig. 4A-D**). For this analysis, there was similar representation of cross links from both models (**Fig. 4E**). We next measured the distances between each pair of cross-linked lysine ε-amino group to investigate if the mapped cross links existed within the expected distance constraint of tPhoX (<35Å). A total of 37 out of 40 XL-peptides were within the distance constraint (**Fig. 4F**). Two cross links that violate this constraint exists either between MYH4 head group and TPM1 or within MYL1 of the Actomyosin Complex. These domains are structurally re-organised following binding of calcium and ATP and remodelling of actin-myosin cross-bridges during muscle contraction (Sweeney & Houdusse, 2010, Irving et al., 2000). Therefore, excluding these examples, 97.5% of cross links across these complexes fall within the distance constraint of tPhoX.

**Figure 4.**
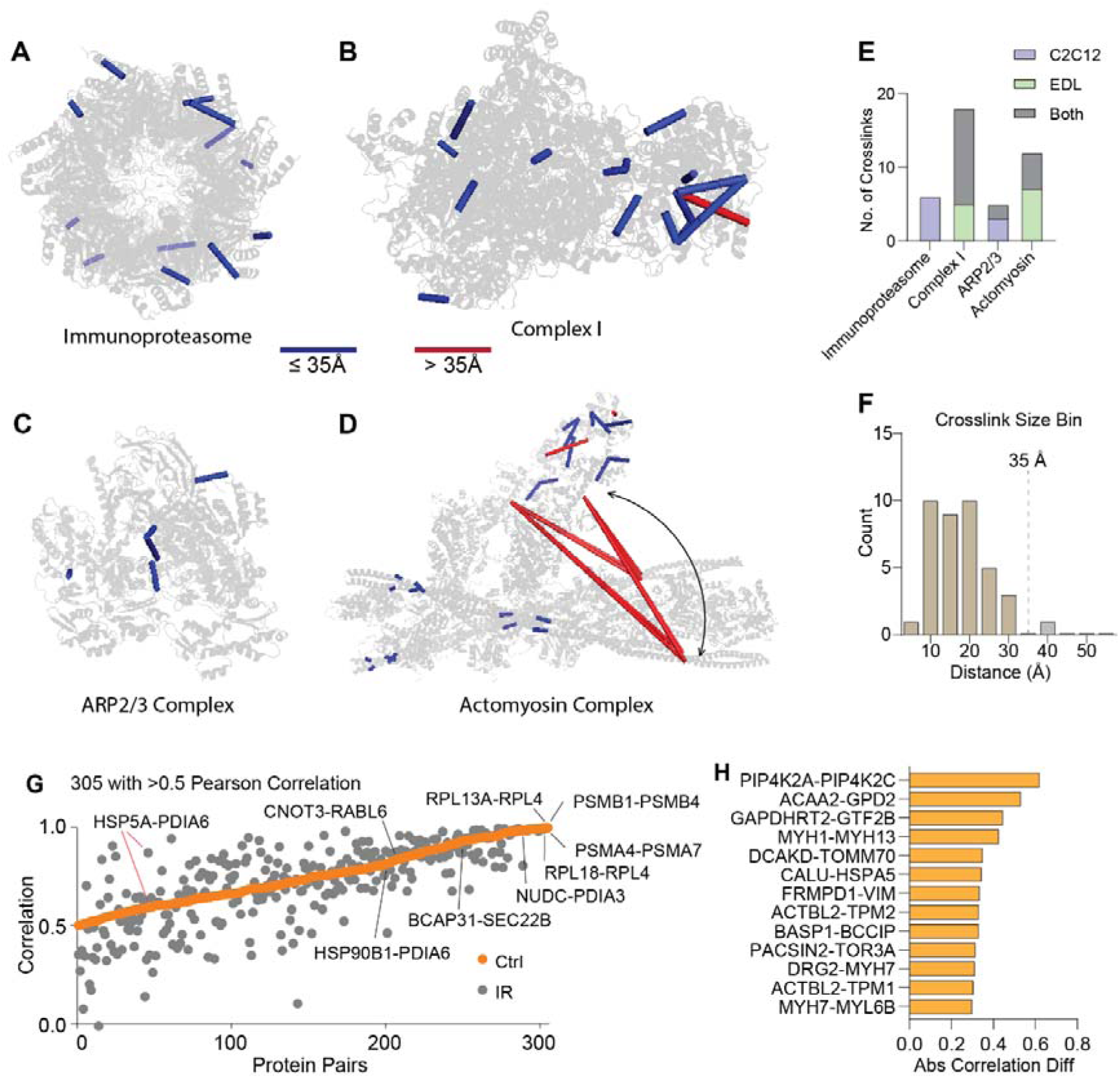
Integration and validation of whole cell cross linking. (**A-D**) Cross linked peptides mapped onto experimentally validated protein complex structures. (**E**) Distribution of mapped cross links and their model of origin. (**F**) Distribution of cross links that fall within the distance constraint of tBu-PhoX. (**G**) Integration of total inter-protein cross links with pairwise correlation analysis of PCP-MS protein elution profiles. (**H**) Cross linked protein pairs with the greatest absolute difference in correlation between healthy and insulin resistant states.

Integration of i) PCP-MS in C2C12, ii) qXL-MS in C2C12, or iii) qXL-MS in EDL muscles identified 435 PPIs in at least two out of the three datasets (**Supplemental Table S8**). Of these, 305 protein-pairs had a correlation >0.5 in the C2C12 PCP-MS and were also identified as cross-linked in the C2C12 and/or EDL qXL-MS data. These interactions include subunits of well-established complexes such as the proteasome and ribosome, but also shed light on novel direct PPIs not annotated in the latest integrated Hu.MAP3db such as, CNOT3-RABL6, BCAP31-SEC22B, PDIA6-HSP90B1, and PDIA6-HSP5A (**Fig. 4G**). The strength of several protein pair co-elution correlations in the PCP-MS data was seen to change with induction of IR, and this was validated as a change in XL-peptide abundance (**Fig. 4H**). For example, an association between ACAA2-GPD2 had both decreases in correlation strength in PCP-MS and decreases in XL-peptides with IR. Both proteins are localised to the mitochondria however the ACAA2 catalyses the breakdown of fatty acids in the beta-oxidation pathway while GPD2 catalyses the conversion of glycerol-3-phosphate into dihydroxyacetone phosphate and indirectly reduces free-glycerol levels essential for fatty acid synthesis. Hence, a decrease in the ACAA2-GPD2 interaction is a fascinating and unexplored area of fatty acid metabolic regulation. As mentioned above, protein processing in the ER also stood out as having dysregulated PPIs identified by both PCP-MS and qXL-MS. A large decrease in the correlation strength was observed between CALU-HSPA5 with IR, and this correlated with a significant decrease in multiple XL-peptides between these two proteins. Both proteins are chaperones localised to the ER and play critical roles in protein folding and translocation but the role of this interaction in IR has, to our knowledge, not been investigated. PDIA6, an ER-resident protein-disulfide isomerase and important for the regulation of protein folding via modulating the oxidation status of cysteines also displayed several dysregulated PPIs identified by both PCP-MS and qXL-MS following the induction of IR. Point mutations in *Pdia6* or loss-of-function cause severe diabetes which has primarily been attributed to a reduction in insulin processing and production (Eletto et al., 2016, Gorasia et al., 2016, Chhabra et al., 2021), however a role of this protein in skeletal muscle insulin sensitivity is unknown. Hence, our quantitative interactome map of skeletal muscle IR provides new potential molecular mechanisms driving changes in metabolic adaptations, protein quality control and function.

### Dysregulated PDIA6 modulates protein redox status and insulin sensitivity

PDIA6 stood out as a central hub dysregulated with IR and therefore, we further investigated its potential contributions to skeletal muscle insulin sensitivity via changes in the PPIs and cysteine oxidation. We identified 18 direct PPIs with PDIA6 including several not previously reported in Hu.MAP3db that were also dysregulated with IR (**Fig. 5A**). A total of 34 unique XL-peptides were identified between PDIA6 and HSPA5 (BIP), a known ER stress chaperone responsible for regulating the UPR, and these were all down-regulated with the induction of IR (**Fig. 5B**). Interestingly, recent phosphoproteomics studies in human skeletal muscle have identified both exercise- and insulin-regulated phosphosites at S129 and S428 on PDIA6 (Blazev et al., 2022, Needham et al., 2022) (**Fig. 5B-C**), and proteomic analysis of muscle biopsies from type-2 diabetic patients have shown significant downregulation of PDIA6 protein abundance (Ohman et al., 2021). Analysis of our PCP-MS data revealed a complex re-organisation of PDIA6 elution profiles in IR C2C12 myotubes with or without acute 30 min insulin stimulation (**Fig. 5D**). IR was associated with a greater distribution of PDIA6 across a range of higher molecular weights (blue arrow). There was also an acute insulin-dependent shifts that differ in the IR state (red arrow). BN-PAGE and western-blotting was performed in control or palmitate-induced IR C2C12 myotubes with or without acute insulin stimulation further supporting complex re-organisation including decreased monomeric forms of PDIA6 and increased associations to heavy molecular weights with IR which were further remodelled with acute insulin stimulation (**Fig. 5E**).

**Figure 5.**
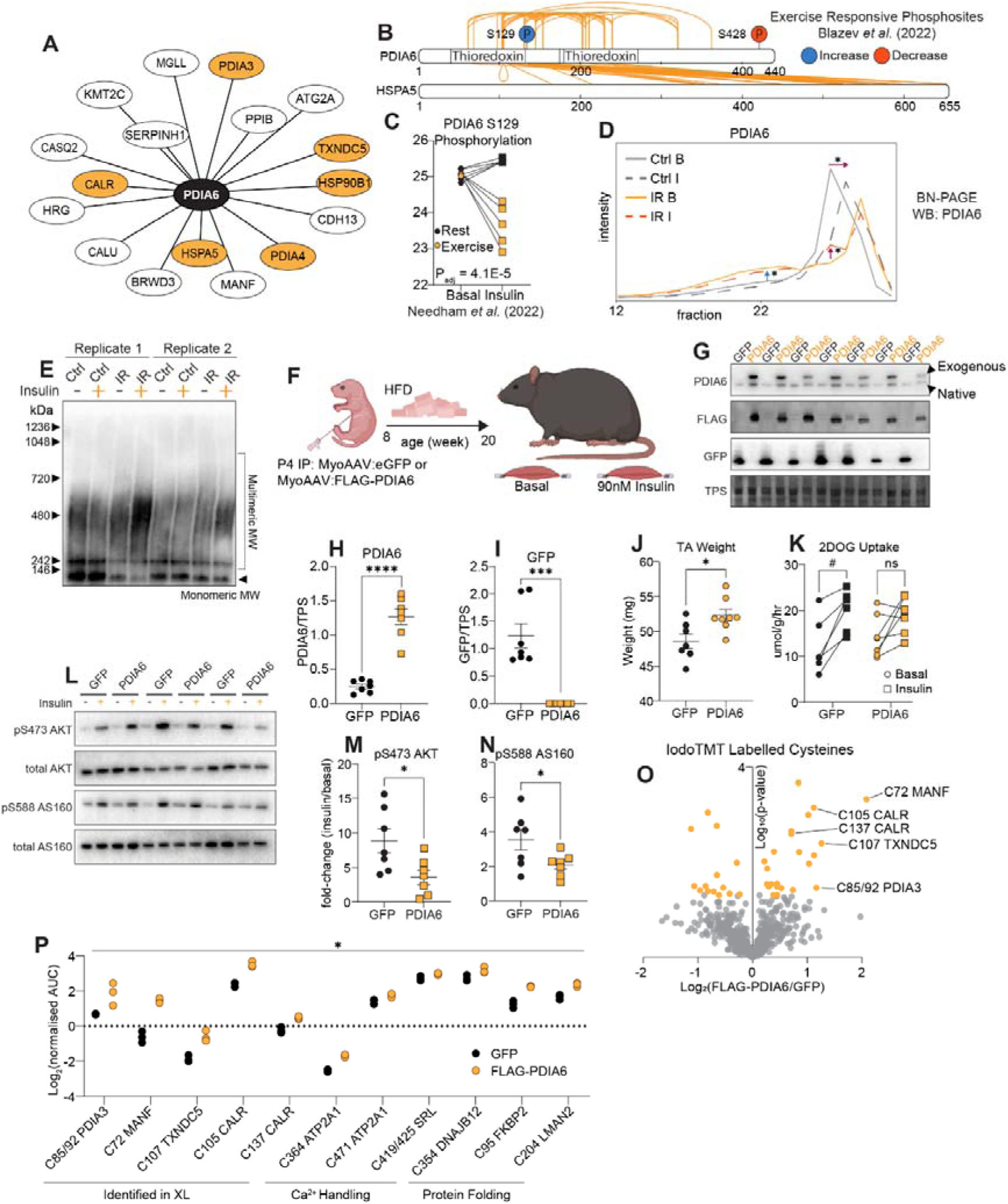
Phenotypic investigation of PDIA6 and its role in skeletal muscle insulin sensitivity. (**A**) Visual schematic detailing functional domains and phosphorylation sites of PDIA6, and quantified cross links between PDIA6 and BIP. (**B**) Cross linked interactome network of PDIA6 where purple lines denote experimentally validated interactions. (**C**) Phosphorylation of S129 on PDIA6 in response to insulin stimulation and exercise treatment (n=5). (**D**) Elution profile of PDIA6 quantified via PCP-MS (n=3). (**E**) Western blotting of PDIA6 on natively lysed healthy and insulin resistant (IR) C2C12 myotubes that were either basal or treated with insulin stimulation separated via BN-PAGE (n=2). (**F**) Experimental timeline investigating FLAG-PDIA6 overexpression and its role in skeletal muscle insulin sensitivity during the development of IR. (**G-I**) Western blotting confirming overexpression of either PDIA6 or GFP. (**J**) *Tibialis anterior* weights and (**K**) *soleus* 2DOG uptake of GFP and FLAG-PDIA6 overexpressing mice. (**L-N**) Western blotting of downstream insulin signalling and its quantification (n=7-8). (**O**) Volcano plot of iodoTMT labelled cysteins in GFP vs FLAG-PDIA6 *tibialis anterior* muscles (n=3). (**P**) Comparison of significantly regulated cysteines on cross linked or functionally relevant proteins. *p<0.05, **p<0.01, ***p<0.001 ****p<0.0001, unpaired t-test. #p<0.05, paired t-test.

To further investigate the role of PDIA6 on insulin sensitivity, we overexpressed either GFP control or FLAG-PDIA6 throughout the musculature of 4-day old male C57BL/6J mice via intraperitoneal injection with MyoAAV (**Fig. 5F**). At 8 weeks of age, mice were placed on a HFD for 12 weeks to induce IR and the hindlimbs dissected for various biochemical assays. Western-blotting confirmed over-expression of PDIA6 by 5x-fold (**Fig. 5G-I**) and this was associated with subtle but significant increase in *Tibialis anterior* muscle mass by ∼5% (**Fig. 5J**). *Ex vivo* insulin-stimulated ^3^H-2-deoxyglucose uptake into *Soleus* muscle revealed a subtle decrease in insulin sensitivity following over-expression of PDIA6 (**Fig. 5K**), and this was associated with a reduction in insulin-stimulated phosphorylation of pS473 on Akt and its substrate pS588 on AS160 (**Fig. 5L-N**). We next aimed to identify specific cysteine residues with altered redox status following the over-expression of PDIA6. To achieve this, we performed cysteine-redox proteomics using isobaric stable isotope labelling with iodoTMT. Here, an aliquot of liquid nitrogen powdered *gastrocnemius* muscles overexpressing GFP or FLAG-PDIA6 were lysed and free thiols alkylated with NEM (n=3 each). Oxidised thiols were reduced and then labelled with iodoTMT 6-plex prior to protein digestion and 2DLC-MS/MS. This quantified 578 cysteine-labelled peptides where 23 and 14 had increased and decreased oxidation, respectively following PDIA6 over-expression (**Fig. 5O**) (**Supplementary Table S9)**. The most significantly oxidised cysteines were on proteins found to be crosslinked with PDIA6 including MANF, CALR, TXNDC5 and PDIA3. Proteins involved in Ca^2+^ handling including ATP2A1 and SRL, and other proteins involved in protein folding and sorting including DNAJB12, FKBP2 and LMAN2 also contained increased oxidised cysteines (**Fig. 5P**). This is consistent with current literature as PDIA6 is known to form condensates where it anchors other folding chaperones, and that formation of these condensates is Ca^2+^ dependent (Leder et al., 2025).

## Discussion

In this study, we performed PCP-MS analysis of healthy or palmitate-induced IR C2C12 myotubes treated with or without acute insulin stimulation to investigate changes in protein interactions. Normalization and differential analysis of these data using CCProfiler revealed proteome wide reorganization of known protein complexes and proteome wide changes in protein elution profiles in response to both IR and insulin stimulation, suggesting global changes in protein interactions. Utilizing the membrane permeable and tPhoX crosslinker, a complementary qXL-MS analysis of healthy and IR C2C12 myotubes treated with or without acute insulin stimulation was performed to orthogonally identify global changes in protein-interactions. This proteome-wide approach was enabled by extensive fractionation and enrichment of crosslinked peptides which revealed changes in the interactome with IR. However, unlike PCP-MS, qXL-MS did not capture changes in PPIs with acute insulin stimulation despite cells being stimulated with insulin for 30 min followed by an additional 30 min of cross-linking in the presence of insulin. The lack of acute changes captured with this approach is surprising. One explanation is that acute changes in PPIs are substoichiometric and only a very small fraction of XL-peptides change in abundance. PCP-MS on the other hand can separate these re-organised PPIs and this orthogonal fractionation improves the ability to detect low abundant changes.

We next established qXL-MS to analyse changes in PPIs in intact tissue; whole EDL muscles from chow- and high-fat-diet mice. As cross-linked proteins are low abundant, extensive enrichment and separation was needed for this analysis. This was made possible using an IMAC enrichable crosslinker (tPhoX) followed by size exclusion chromatography to enrich larger XL-peptides, and multiple rounds of fractionation via subcellular and reversed-phased chromatography. Mapping of selected cross links to experimentally validated protein complex structures revealed that 95% fell within distance constraints of tPhoX, with violations only occurring in known flexible regions. We identified more XL-peptides in cell culture compared to intact muscle which may be attributed to the greater ability of tPhoX to permeate cells rather than whole tissue. Pathway enrichment analysis revealed greater capture of interacting proteins associated with contractile function, calcium handling and glycolysis in the *EDL* muscles. On the other hand, cross linking of C2C12 myotubes captured protein-protein interactions associated with vesicular transport, fatty acid biosynthesis, oxidative phosphorylation, thermogenesis and protein processing in the ER. The different subsets of proteins captured may be due to proteomic differences with the whole tissue being dominated by contractile proteins and ultimately suggests a complementary relationship between the data generated from both models, highlighting the value of each approach.

One of the major pathways captured was protein processing in the ER and a potential link to the unfolded protein response (UPR) which has been associated with IR. Oℒzcan et al. (2004) showed that ER stress results in insulin resistance via a suppression of insulin receptor signalling. This occurred through IRE1α-dependent activation of JNK which led to inhibition of IRS-1 and therefore downstream signalling. These findings, however, were predominantly observed in liver and adipose tissues instead of muscle. Brown et al. (2020) showed that ER stress and therefore activation of the UPR inhibits insulin signalling via depletion of the insulin receptor in C2C12 myotubes. Furthermore, increased markers of ER stress have been repeatedly identified in either obese or insulin resistant muscle tissue (Deldicque et al., 2010, Rieusset et al., 2012) where ER stress induced upregulation of TRB3 has been shown to be responsible for deactivation of AKT and mTOR, and has therefore been implicated as a potential mechanism linking ER stress and insulin resistance in skeletal muscle (Koh et al., 2013, Koh et al., 2006). In our data specifically, there was a general downregulation of PPIs associated with protein handling (PDIA3, RPN1, PDIA1), chaperone activity (HYOU1, CANX, HSPs, BAG2) and proteolysis/ERAD (CAPN1, OS9). This suggests a reduced capacity for these myotubes to alleviate ER stress which may therefore be an underlying cause of the observed IR. This is further supported by accompanying decreases in protein features observed with the insulin receptor, AMPK and several of their downstream proteins. One of the striking interactions within this pathway was the reduced interaction between PDIA6 and BiP. Integration with the PCP-MS analysis revealed coelution of BiP and PDIA6 when separated via BN-PAGE, and that their coelution significantly shifted with induction of IR and with insulin stimulation. Here, IR and acute insulin stimulation induced a shift of PDIA6 away from its monomeric molecular weight and is consistent with how PDIA6 colocalizes and responds to ER stress. Leder et al. (2025) demonstrated that PDIA6 exists in condensates on the ER lumen in homeostatic conditions to limit protein folding to specific substrates only. However, during conditions of ER stress, PDIA6 was observed to disperse throughout the ER where the authors hypothesised that this increased folding capacity at the cost of specificity (Leder et al., 2025). The shift of PDIA6 towards protein complexing may be indicative of this dispersed state in response to IR and insulin stimulation associated ER stress. Interestingly, both the presence of PDIA6 and its ability to dimerize via its a^0^ domain is required for the formation of these condensates where PDIA6 also serves as an anchor to recruit other chaperones (BIP, PDIs, GRP94/Endoplasmin). These events were captured in our dataset where several cross links were identified among the a^0^ domain of PDIA6 however it is difficult to distinguish between intra-protein cross links of PDIA6 and inter-protein cross links between two copies of PDIA6. There were cross links between PDIA6 and known colocalizing chaperones (BIP, PDIA3, PDIA4, CALR) as well as novel chaperones (PPIB, SERPINH1, TXNDC5, CALU) (Leder et al., 2025, Lee et al., 2025). All these cross links were significantly downregulated, further indicating dispersion of PDIA6 condensates.

We therefore hypothesised that a reduction in PDIA6 PPIs leads to reduced protein folding capacity which contributes to ER stress associated with the development of IR, and that increasing PDIA6 may improve protein folding and insulin sensitivity. Therefore, we over-expressed either GFP or PDIA6 throughout the musculature via IP delivery using MyoAAV. These mice were then placed on a high fat diet for 12 weeks to induce IR. Contrary to expectations, over-expression of PDIA6 resulted in reduced skeletal muscle insulin sensitivity. Eletto et al. (2014) has shown that PDIA6 acts to limit IRE1α activity and that PDIA6 deficient cells hyper respond to ER stress in C. elegans (Eletto et al., 2014). Overexpression of PDIA6 may therefore blunt insulin signalling via suppression of the UPR. Early work by Kangdong et al. (2024) revealed that PDIA6 stabilizes TRAF4 to promote AKT1/mTOR/p70S6K signalling in esophageal squamous cell carcinoma cells (Liu et al., 2024). Overexpression of PDIA6 may therefore drive chronic activation of this pathway and therefore drive negative feedback loops that attenuate insulin signalling, resulting in the observed blunting of insulin-stimulated glucose uptake (Rozengurt et al., 2014).

To investigate potential mechanisms, we performed global cysteine redox proteomics and show that over-expression of PDIA6 in IR skeletal muscle increased cysteine oxidation of crosslinked partners (MANF, CALR, TXNDC5, PDIA3). These proteins are ER-resident chaperones, and notably, altered MANF expression in metabolic tissues has been linked to regulation of metabolic disease (Tang et al., 2022). Additional changes in cysteine oxidation were observed in other protein-folding chaperones (DNAJB12, FKBP2, LMAN2) as well as Ca²⁺-handling proteins (ATP2A1, SRL), consistent with PDIA6 co-existing in an ER-resident chaperone network. Formation of PDIA6-associated condensates is Ca²⁺-dependent. Thapsigargin, which blocks ATP2A1 activity, rapidly depletes ER Ca²⁺, induces ER stress, and triggers phase separation of these PDIA6-containing droplets. This dependency is mediated through three distinct Ca²⁺-binding sites on PDIA6 (Leder et al., 2025). Together, these data suggest that PDIA6-dependent modulation of disulfide bond formation in ATP2A1 may allosterically tune its activity, establishing a potential Ca²⁺-sensitive feedback mechanism linking protein folding stress to Ca²⁺ handling. (Leder et al., 2025).

### Limitations & Future Directions

While XL-MS yields many advantages including the ability to covalently trap PPIs within a cell, the application *in vivo* is still limited by the cost and feasibility of intravenous infusion of large quantities of cross linker. We therefore performed our experiments *ex vivo* where *EDL* muscles were freshly excised from mice prior to 2 hours of cross linking in physiological conditions. The effects of this *ex vivo* procedure on the interactome are unknown and serves as a primary limitation of this data. Furthermore, while it was necessary to maximize depth of analysis, it is also unclear how the application of several orthogonal methods of fractionation have on the proteins captured and their quantitation. Additionally, it is still difficult to distinguish intra-protein cross links from inter-protein cross links across homomeric complexes unless the two XL-peptides have the identical peptide sequences. Hence, experiments utilising simultaneous XL-MS and PCP-MS may provide additional benefits which has previously been used to study mitochondria isolations (Hevler et al., 2021). Future experiments are required to elucidate the exact role of PDIA6 in the context of metabolic disease. For example, it would be interesting to investigate if Ca^2+^-dependent PDIA6 condensates change in response to metabolic stress.

## Conclusion

This study was able to demonstrate the ability for XL-MS to capture protein interactions on a proteome wide scale. Bolstered by the use of extensive fractionation to increase depth and multiplexing to boost sensitivity, we were able to quantify changes in direct protein interactions in healthy and IR muscle cells, and for the first time whole skeletal muscle tissue marking a new milestone in the applications of XL-MS. Integration with protein co-elution data provided a high confidence quantitative interactome that highlighted protein interactions associated with the development of skeletal muscle insulin sensitivity. This included the ER chaperone PDIA6 where its overexpression negatively impacted insulin stimulated glucose uptake while promoting disulfide bond formation in key regions of its interacting proteins.

## Acknowledgements

We thank Nicholas Williamson, Ching-Seng Ang, Shuai Nie, Swati Varshney and Michael Leeming for instrument support in the Bio21 Mass Spectrometry and Proteomics Facility. We also thank Laura Dagley and Steve Binos for instrument support in the Walter and Eliza Hall Proteomics Facility. Finally, we thank Pin-Lian Jiang, Fan Liu, Rosa Viner and Ryan Bomgarden for advice on tPhoX cross-linking and TMT acquisition protocols. This research was supported by access to the Melbourne Mouse Metabolic Phenotyping Platform at The University of Melbourne. This work was funded by a University of Melbourne Driving Research Momentum Grant, and an NHMRC Emerging Leader Investigator Grant APP2009642 and Australian Research Council Discovery Grant DP250100201 to B.L.P.

## Methods

### Mouse housing

All mouse experiments were approved by The University of Melbourne Animal Ethics Committee (AEC ID1914940) and conformed to the National Health and Medical Research Council of Australia guidelines regarding the care and use of experimental animals. C57BL/6J mice (JAX 000664) were obtained from Animal Resource Centre (WA, Australia). Mice were housed at 22°C (+/-1°C) in groups of five/cage and maintained on a Standard Chow diet (Specialty Feeds, Australia) or a High Fat Diet (Specialty Feeds Australia, #SF04-001) with a 12-hour light/dark cycle and ad libitum access to food and water.

### Insulin stimulated 2-deoxyglucose uptake

Mice were anesthetized with isoflurane (4% in oxygen at 1 l/min and transferred to a dissecting microscope stage with isoflurane inhalation nose piece (2% in oxygen at 1L/min). Depth of anaesthesia was assessed via the lack of leg and optical reflexes for at least one minute to ensure head position did not affect normal breathing. After confirming anaesthesia, skin from the hind legs was removed, and soleus muscles were sutured using 4.0 braided suture at both proximal and distal ends at the tendomuscular junction. Muscles were excised and incubated in Modified Krebs Buffer (MKB) (116 mM NaCl, 4.6 mM KCl, 1.16 mM KH2PO4, 25.3 mM NaHCO3, 2.5 mM CaCl2, 1.16 mM MgSO4) containing 0.1% BSA, 2 mM Pyruvate and 7 mM cold-mannitol in a myograph (DMT, Denmark; #820MS) at 30°C with constant gentle bubbling of 5% medical carbon dioxide in oxygen. Left and right soleus muscles were adjusted to resting tension (∼5 mN) and allowed to equilibrate for 10 min before being stimulated (left leg no insulin, right leg 90 nM insulin) for 30 min. During the last 10 min of stimulation, the buffer was replaced with MKB containing 0.375 μCi/ml 2-deoxy-d-[1,2-3H] glucose and 0.05 μCi/ml d-[14C] mannitol after which muscles were immediately immersed in ice-cold PBS to stop further glucose uptake. Each muscle was then excised of its sutures/tendons and frozen in liquid nitrogen for future lysis in 400 μl of 2% SDS via tip-probe sonication (QSonica). Measurement of radiolabelled 2-deoxyglucose was carried out in a Tri-Carb 4910 TR liquid scintillation analyzer (Perkin Elmer) by adding 200 μl of muscle lysate to 4ml scintillation fluid (Ultima Gold, Perkin Elmer). Glucose uptake rates were calculated as described in (Kjobsted et al., 2021) where significance was determined via paired t-test.

### Western-blotting

Protein was separated on NuPAGE 4-12% Bis-Tris protein gels (ThermoFisher Scientific) in MOPS SDS Running Buffer at 160 V for 1 h at room temperature. The protein was transferred to PVDF membranes (Millipore; #IPFL00010) in NuPAGE Transfer Buffer at 20 V for 1 h at room temperature and blocked with 5% skim milk in Tris-buffered saline containing 0.1% Tween-20 (TBST) for at least 30 min at room temperature with gentle shaking. The membranes were incubated overnight in primary antibody with 5% BSA in TBST with gentle shaking at 4°C and washed three times in TBST at room temperature. The membranes were incubated with HRP-secondary antibody in 5% skim milk in TBST for 45 min at room temperature and washed three times with TBST. Protein was visualized with Immobilon Western Chemiluminescent HRP Substrate (Millipore; #WBKLS0500) and imaged on a ChemiDoc (BioRad). Densitometry was performed in ImageJ (Schneider et al., 2012). Total protein staining was performed with BLOT-FastStain (G-Biosciences; #786-34) where membranes were incubated in Fixer solution (diluted 1:10) for 2-3 minutes with gentle shaking followed by incubation in Developer solution (diluted 1:4) for 30 minutes with gentle shaking. Staining was imaged on a ChemiDoc (BioRad).

### SYPRO Staining

For in-gel total protein staining of samples separated via SDS-Page, SYPRO Ruby Protein Gel Stain was used (Sigma-Aldrich; S4942). Gels were fixed in 40% methanol, 10% acetic acid for 30 minutes with gentle shaking at room temperature. Gels were then incubated in approximately 50ml of SYPRO stain and incubated overnight with gentle shaking at room temperature. Gels were then washed 3 x 5mins in milliQ and imaged on a ChemiDoc (BioRad).

### Cell Culture of C2C12 myotubes and induction of insulin resistance

C2C12 cells were grown in Dulbecco’s Modified Eagle Medium (DMEM) (GIBCO by Life Technologies; # 11995065), supplemented with 10% fetal bovine serum (FBS) (Life Technologies; #26140079), pyruvate and GlutaMAX (GIBCO by Life Technologies). Cells were kept at 37°C and 5% CO2 in a humidified incubator Direct Heat CO2 Incubator featuring Oxygen Control (In Vitro Technologies). At 90% confluence, the cells were differentiated into myotubes with the replacement of 10% FBS with 2% Horse Serum and cultured for a further 3 days. Myotubes were then incubated in 1ml of DMEM, 2% Horse Serum, Pyruvate, GlutaMAX with 0.2% BSA (fatty acid free) with or without 500µM palmitic acid for 24 hours to induce insulin resistance (IR). For IR incubation media, 5 mM palmitic acid dissolved in ethanol was conjugated with 2% BSA in DMEM, Pyruvate and GlutaMAX (GIBCO by Life Technologies) and diluted 10-fold. IR incubation media was then filter sterilised prior to incubation. On the day of harvest, cells were incubated in 1ml of DMEM, Pyruvate, GlutaMAX with 0.2% BSA with or without 500µM palmitic acid for 2 hours to achieve a basal state. Relevant cells were then stimulated with 100nM insulin for 30 minutes. Cells were washed twice with cold PBS, lysed in 100ul of native lysis buffer (0.25% Triton-X100, 10mM Tris.HCl, 50mM NaCl, 1mM TCEP, 1mM EDTA, EDTA-free PIC, phosIC, pH 7.5) and transferred to cold 1.5ml tubes and left on ice for 10 minutes. Cells were gently mixed with a pipette, spun down at 20,000 g for 20 minutes at 4°C, after which proteins were quantified with Qubit and normalized for separation via Blue Native-PAGE and SDS -PAGE.

### Blue Native-PAGE Separation for PCP-MS

Native lysates were separated on a NuPAGE 3-8% Tris-Acetate gel (Invitrogen; #EA0375BOX) at 150V for 2 hours with Tris-Glycine Native running buffer (Invitrogen; LC2672). Coomassie staining was then performed to visualize gel lanes. Briefly, gels were fixed in 40% methanol, 10% acetic acid for 10 minutes and directly incubated in 50ml of QC Colloidal Coomassie stain (BioRad; #1610803) overnight with gentle shaking at room temperature. Staining solution was removed, and gel was washed with milliQ overnight. Each gel lane was cut into 30 x 1.5mm fractions and each fraction was transferred into individual wells in a 96-well plate.

Gel fractions were washed twice with 1ml of 40% acetonitrile in 50mM ammonium bicarbonate at 1500 x rpm for destaining. 500ul of acetonitrile was added and plate was shaken at 1500 x rpm to dehydrate gel fractions. Acetonitrile was removed and gel fractions were then rehydrated with 40ul of trypsin (13ng/ul; Sigma; #T6567) at 4°C for 1.5 hours and excess removed. Gels were then covered with 50ul of 50mM ammonium bicarbonate to prevent dehydration then incubated overnight at 37°C.

The next day, 50ul of 20mM TCEP (Sigma; #75259) and 80mM CAA (Sigma; #22790) in 100mM Tris pH8.5 was added to each well and heated at 37°C for 30 minutes for reduction and alkylation. 100ul of 1% TFA was added and mixed for 15 minutes at 100 x rpm at room temperature and the flowthrough was collected. Gel pieces were washed again with 100ul of 80% acetonitrile 0.1% TFA for 15 minutes at 100 x rpm at room temperature after which both flowthroughs containing digested peptides were purified with in-house packed SDB-RPS (Sigma; #66886-U) microcolumns. The purified peptides were resuspended in 2% acetonitrile in 0.1% TFA and stored at -80°C prior to direct injection by LC-MS/MS (Harney & Larance, 2023).

### Cross linking-mass spectrometry analysis of C2C12 myotubes and *EDL* muscles *ex vivo*

C2C12 cells were grown in Dulbecco’s Modified Eagle Medium (DMEM) (GIBCO by Life Technologies; # 11995065), supplemented with 10% fetal bovine serum (FBS) (Life Technologies; #26140079), pyruvate and GlutaMAX (GIBCO by Life Technologies). Cells were kept at 37°C and 5% CO2 in a humidified incubator Direct Heat CO2 Incubator featuring Oxygen Control (In Vitro Technologies). At 90% confluence, the cells were differentiated into myotubes with the replacement of 10% FBS with 2% Horse Serum and cultured for a further 3 days. Myotubes were then incubated in 1ml of DMEM, 2% Horse Serum, Pyruvate, GlutaMAX with 0.2% BSA (fatty acid free) with or without 500µM palmitic acid for 24 hours to induce insulin resistance (IR). For IR incubation media, 5 mM palmitic acid dissolved in ethanol was conjugated with 2% BSA in DMEM, Pyruvate and GlutaMAX (GIBCO by Life Technologies) and diluted 10-fold. IR incubation media was then filter sterilised prior to incubation. On the day of harvest, cells were incubated in 1ml of DMEM, Pyruvate, GlutaMAX with 0.2% BSA with or without 500µM palmitic acid for 2 hours to achieve a basal state. Relevant cells were then stimulated with 100nM insulin for 30 minutes. Media was removed and cells were washed in cold PBS. Myotubes were then scraped in 10 ml of cold PBS, transferred to a 15 ml falcon tube and spun down at 150 g for 10 minutes at 4 degrees. A stock solution of 50 mM of t-butyl-PhoX (tBu-PhoX) (Thermo Scientific, A52287) resuspended in DMSO was gently heated at 37 degrees Celsius until fully dissolved. Stock solution was then diluted with PBS to 2 mM for tBu-PhoX incubation solution. PBS was removed then removed from the myotubes where the cell pellet was resuspended in 500 µl of PBS containing 2 mM tBu-PhoX and incubated for 30 minutes at room temperature. After incubation, cells were quenched with 10 µl of 1 M Tris pH 8.5 and gently mixed on a shaker for 5 minutes. 10 ml of cold PBS was then added to each sample, cells were spun down at 150 g for 10 minutes at 4 degrees Celsius after which PBS was removed and samples proceeded with lysis for subcellular fractionation.

*EDL* muscles were excised and incubated in 10 mM of tBu-PhoX in 1 ml of MKB (116 mM NaCl, 4.6 mM KCl, 1.16 mM KH2PO4, 25.3 mM NaHCO3, 2.5 mM CaCl2, 1.16 mM MgSO4, 2 mM pyruvate, 11mM glucose) for 2 hours at 30°C with constant gentle bubbling of 5% medical carbon dioxide in oxygen. A stock solution of 200 mM of tBu-PhoX resuspended in DMSO was gently heated at 37 degrees Celsius until fully dissolved where 50 µl was added to 950 µl of MKB for a final concentration of 10 mM tBu-PhoX. It is important to exclude BSA from the incubation buffer as tBu-PhoX will start cross-linking any proteins present. After incubation, *soleus* muscles were quenched in ice cold 100 mM Tris pH7.4 buffer for 5 minutes on ice and snap frozen for lysis with subcellular fractionation.

### Subcellular Fractionation

Subcellular fractionation of C2C12 myotubes and *EDL* muscles was performed using a Subcellular Fractionation Kit (Thermo Scientific; 87790) according to their product protocol. Briefly, proprietary cytoplasmic (CEB), membrane (MEB), nuclear (NEB) and pellet (PEB) extraction buffers were prepared by adding 1:100 of 100X Halt Protease Inhibitor Cocktail to each extraction buffer immediately before use. Recommended volumes of each buffer were used based on the starting material. An additional nuclear extraction buffer (NEB+) was prepared by adding 5µL of 100mM CaCl_2_ and 3µL of Micrococcal Nuclease per 100µL of NEB to target extraction of chromatin bound.

For C2C12 myotubes, cross-linked cells were centrifuged at 500 g for 3 minutes after which the supernatant was carefully removed and discarded, leaving the cell pellet as dry as possible. Ice-cold CEB containing protease inhibitors was added to the cell pellet and incubated at 4 degrees Celsius for 10 minutes with gentle mixing. Cell lysate was then centrifuged at 500 g for 5 minutes where supernatant (cytoplasmic extract) was transferred to a clean pre-chilled tube on ice. Pellet was then subject to further fractionation.

For *EDL* muscles, snap frozen cross-linked tissues were added to ice-cold CEB containing protease inhibitors and homogenize using a Polytron handheld homogenizer on speed 2 for ∼20 seconds. Homogenized tissue was then transferred to a Pierce Tissue Strainer attached to a 15mL conical tube and centrifuged at 500 g for 5 minutes. Supernatant (cytoplasmic extract) was then transferred to a clean prechilled tube while the pellet was subject to further fractionation.

Ice-cold MEB containing protease inhibitors was added to sample pellets and vortexed for 5 seconds on the highest setting. Lysate was incubated for 10 minutes at 4 degrees Celsius with gentle mixing. Lysates were then centrifuged at 3000 g for 5 minutes and supernatant (membrane extract) was transferred into a clean pre-chilled tube on ice.

Ice-cold NEB containing protease inhibitors was added to the pellet and vortexed for 15 seconds on the highest setting. Lysate was incubated for 30 minutes at 4 degrees Celsius with gentle mixing. Lysates were then centrifuged at 5000 g for 5 minutes and supernatant (soluble nuclear extract) was transferred into a clean pre-chilled tube on ice.

Ice-cold NEB+ containing protease inhibitors, CaCl_2_ and Micrococcal Nuclease was added to the pellet and vortexed for 15 seconds on the highest setting. Lysate was incubated for 15 minutes at 37 degrees Celsius in a water bath. Vortex lysates again on the highest setting for 15 seconds and centrifuge at 16,000 g for 5 minutes. Supernatant (chromatin-bound nuclear extract) was then pooled with the soluble nuclear extract due to limited yield.

Remaining cell pellet was then resuspended in 200 µl of Gdm lysis buffer (6 M Gdm, 10 mM TCEP (Sigma; #75259), 40 mM CAA (Sigma; #22790) in 100 mM Tris pH 8.5) (cytoskeletal fraction) and boiled at 95 degrees Celsius for 10 minutes for reduction and alkylation, after which all fractions proceeded with further processing.

10 mM TCEP (Sigma; #75259) and 40 mM CAA (Sigma; #22790) was added to the cytoplasmic, membrane and pooled nuclear fractions which were then reduced and alkylated at 45 degrees for 5 minutes in a water bath. The cytoskeletal fraction was then diluted 1:1 with water after which all proteins were extracted using a methanol-chloroform precipitation method.

### Methanol-Chloroform Protein Precipitation

Briefly, 4 volumes of cold methanol, 1 volume of cold chloroform, 3 volumes of cold LC-MS water were added to each fraction in that order with mixing in between. Fractions were spun down at 4,500 g for 5 minutes after which the upper phase was carefully removed and discarded, being careful not to disturb the protein precipitate ‘disk’ at the water: chloroform interface. 3 volumes of Methanol were added, fractions were vortexed and centrifuged for 4,500 g for 5 minutes. All solution was then removed as proteins were pelleted completely. 5 volumes of cold methanol were added, fractions were vortexed again and centrifuged for 4,500 g for 5 minutes. All remaining solution was removed, and protein pellet was left to dry at room temperature. All fractions were then resuspended in 100 µl of Digestion Buffer (10% 2,2,2-Trifluoroethanol (Sigma; #96924)) in 100 mM HEPEs pH 8.5). Protein was quantified with BCA (ThermoFisher Scientific) and the lowest amount of protein across all samples present in each subcellular fraction was normalized to a final volume of 100 µl in Digestion Buffer, and digested with sequencing grade trypsin (Sigma; #T6567) and sequencing grade LysC (Wako; #129-02541) at a 1:50 enzyme:substrate ratio overnight at 37°C with shaking at 2000 x rpm.

### TMT Labelling and cross-linked peptide enrichment

All XL-MS samples and their resulting fractions were labelled with isobaric tags for quantification. Every 1 µg of peptide was labelled with 2 µg of 16-plex tandem mass tags (TMTpro) in 50% acetonitrile at room temperature for 1.5 h (Zecha et al., 2019). The reaction was deacylated with 0.3% (w/v) of hydroxylamine for 10 min at room temperature and quenched to a final volume of 0.1% formic acid (FA) and <5% acetonitrile. At this point, the 16 TMT labelled samples per experiment were pooled. A total of 4 TMT 16-plex experiments were performed where each experiment consisted of a single fraction from 16 samples (8x HFD and 8x Chow mice or 8x IR and 8x Control C2C12 myotubes).

Pooled fractions (4 total) were then loaded onto a HLB-SPE column (Waters Corp; #186008055) pre-conditioned with 1 ml of methanol followed by 1 ml of acetonitrile and finally 2 ml of 0.1% FA. Columns were washed with 2 ml of 0.1% FA and eluted with 1 ml of 80% acetonitrile, 0.1% FA. Phosphopeptides were pre-cleared with 10ul of Fe-NTA Magnetic Agarose beads (Thermo Scientific; A52284) per 500 µg of peptide pre-washed with 1 ml of 80% acetonitrile, 0.1% FA. The unbound flowthrough which contained our protected cross-linked and regular peptides were collected and dried down in a Speedvac at 45 degrees Celsius. The cross-linked/regular peptide flowthrough was resuspended in 100 µl of 0.5% TFA and bath sonicated for 5 minutes. Samples were then shaken at room temperature for 2 hours at 1,800 RPM which deprotected cross-linked peptides. Each sample was diluted to a final concentration of 80% acetonitrile and 0.1% TFA with 400 µl of acetonitrile and transferred to another lo bind tube containing 10ul of Fe-NTA Magnetic Agarose beads (Thermo Scientific; A52284) per 500 µg of peptide pre-washed with 1 ml of 80% acetonitrile, 0.1% TFA. Samples were shaken again for 30 minutes at room temperature at 1,800 RPM to bind deprotected PhoX-modified peptides, after which the unbound flowthrough containing regular peptides only (total proteome) was collected and saved. Beads were washed with 1 ml 80% acetonitrile and 0.1% TFA three times and saved. PhoX-modified peptides were then eluted off the beads twice with 100 µl of 5% ammonium hydroxide shaken at 2000 RPM for 2 minutes at room temperature. The PhoX-modified peptide eluates, as well as regular non-modified peptide fractions for global total proteomic analysis were then loaded onto in-house packed C8 microcolumns to remove any trace agrose beads (3M Empore; #11913614) and centrifuged at 1,000 g and collected in PCR strip tubes. C8 microcolumns were washed with 5 µl of 50% acetonitrile and collected into the same tube to ensure no remaining peptides in the column. Samples were then dried to approximately 50 µl in a Speedvac at 45°C and 200 µl of 99% isopropanol, 1% TFA was added. Peptides were purified by in-house packed SDB-RPS (Sigma; #66886-U) microcolumns and dried by vacuum centrifugation. The PhoX-modified peptides were resuspended in 30% acetonitrile, 0.1% TFA while the non-modified peptides were resuspended in 2% acetonitrile in 0.1% TFA and all samples were stored at -80°C. PhoX-modified peptides were subjected to size exclusion chromatography (SEC) separation as previously described (Harney & Larance, 2023) and 2-3 fractions further separated by by neutral phase C18BEH HPLC into 12 fractions as previously described (Le Couteur et al., 2021) while the regular non-modified peptides were subjected to only neutral phase C18BEH HPLC for separation in 12 fractions.

### IodoTMT Labelling

Global quantification of cysteine oxidation was performed using an iodoTMT isobaric labelling kit (Thermo Scientific; #90101) according to their product protocol. Briefly, *gastrocnemius* muscles over-expressing GFP or FLAG-PDIA6 (n=3 each) were powdered under liquid nitrogen and lysed in 2% SDS in 100mM HEPES pH 6.9 containing 150mM NaCl, 100mM NEM and 5mM EDTA via tip probe sonication. NEM labelling was performed for 1 hour in the dark after which methanol-chloroform precipitation was performed to remove NEM. Protein was then normalised and reduced with 2% SDS in 100mM HEPES pH 6.9 containing 150mM NaCl and 50mM TCEP for 30 mins at 45 °C. Methanol-chloroform precipitation was performed to remove TCEP where protein was normalised again. IodoTMT reagents were added to each sample where labelling occurred for 1 hour at 37 °C in the dark. The reaction was quenched with 20mM DTT for 15 minutes at 37 °C in the dark after which samples were combined. A final chloroform methanol-precipitation was performed, and protein pellet was resuspended in 1.2M Urea in 100mM HEPES pH7.5. Proteins were digested with 1:50 ratio of trypsin and LysC overnight at 37 °C in the dark shaking at 2000RPM. Peptides were acidified to 0.1% TFA and loaded onto a HLB-SPE column for cleanup then lyophilized via a vacuum concentrator. The purified peptides were resuspended in 2% acetonitrile in 0.1% TFA and separated into 12 fractions by neutral phase C18BEH HPLC as described above.

### LC/MS-MS acquisition of PCP-MS, XL-MS and IodoTMT labelled peptides

For XL-MS, peptides were analysed on a Dionex 3500 nanoHPLC, coupled to an Orbitrap Eclipse mass spectrometer (ThermoFischer Scientific) via electrospray ionization in positive mode with 1.9 kV at 275 °C and RF set to 30%. Separation was achieved on a 50 cm × 75 µm column packed with C18AQ (1.9 µm; Dr Maisch, Ammerbuch, Germany) (PepSep, Marslev, Denmark) over 20 or 60 min depending on the offline fractionation UV trace at a flow rate of 300 nL/min. The peptides were eluted over a linear gradient of 3–40% Buffer B (Buffer A: 0.1% v/v formic acid; Buffer B: 80% v/v acetonitrile, 0.1% v/v FA) and the column was maintained at 50 °C. The instrument was operated in data-dependent acquisition mode with an MS1 spectrum acquired over the mass range 350–2000 m/z (60,000 resolution, 200% automatic gain control (AGC) and 50 ms maximum injection time) followed by MS/MS via HCD fragmentation with a cycle time of 3 seconds where only peptides with charge state 3-8 were selected with 1.2 m/z isolation window (30,000 resolution, stepped collision energy of 35±8%, 200% AGC, maximum injection time set to 110 ms, enhanced TMTPro resolution mode).

For total proteomics, data was acquired as above with the following changes. MS1 spectrum were acquired over the mass range 350-1550 m/z followed by MS/MS where only peptides with a charge state of 2-7 were selected with an isolation window of 0.7 m/z (collision energy of 35, maximum injection time set to 54 ms)

For analysis of PCP-MS, peptides were analyzed on a Dionex 3500 nanoHPLC, coupled to an Orbitrap Ascend mass spectrometer (ThermoFischer Scientific) via electrospray ionization in positive mode with 1.9 kV at 275 °C and RF set to 40%. Separation was achieved on a 50 cm × 75 µm column packed with C18AQ (1.9 µm; Dr Maisch, Ammerbuch, Germany) (PepSep, Marslev, Denmark) over 35min at a flow rate of 300 nL/min. The peptides were eluted over a linear gradient of 3–40% Buffer B (Buffer A: 0.1% v/v formic acid; Buffer B: 80% v/v acetonitrile, 0.1% v/v FA) and the column was maintained at 50 °C. The instrument was operated in data-independent acquisition mode with an MS1 spectrum acquired over the mass range 400-1000 m/z (60,000 resolution, 250% automatic gain control (AGC) and 50 ms maximum injection time) followed by MS/MS via HCD fragmentation with 24 windows with 25 m/z isolation and a 1 m/z overlap (30,000 resolution, 2000% AGC and maximum injection time set to auto).

For analysis of IodoTMT labelled samples, peptides were analyzed on a Vanquish Neo nanoHPLC, coupled to an Orbitrap Astral mass spectrometer (ThermoFischer Scientific) via electrospray ionization in positive mode with 1.9 kV at 290 °C and RF set to 40%. Separation was achieved on a 5 cm × 75 µm uPAC column (Thermo Scientific) over 17min at a flow rate of 750 nL/min. The peptides were eluted over a linear gradient of 3–40% Buffer B (Buffer A: 0.1% v/v formic acid; Buffer B: 80% v/v acetonitrile, 0.1% v/v FA) and the column was maintained at 50 °C. The instrument was operated in data-dependent acquisition mode with an MS1 spectrum acquired over the mass range 350–1500 m/z (120,000 resolution, 300% automatic gain control (AGC) and 50 ms maximum injection time) followed by MS/MS via HCD fragmentation with a cycle time of 1 seconds where only peptides with charge state 2-7 were selected with 0.5 m/z isolation window (collision energy of 35%, 100% AGC, maximum injection time set to 20 ms).

### LC/MS-MS data processing of PCP-MS, XL-MS and IodoTMT labelled peptides

Total proteomic data were searched against an in-house generated mouse skeletal muscle specific database containing 11,028 protein entries with SequestHT within Proteome Discoverer v2.5.0.4 ( 24226387). The database was created by filtering the mouse UniProt database (September 2019) with any protein identified in the following proteomic, phosphoproteomic, glycoproteomic or ubiquitinomic datasets (Deshmukh et al., 2015, Goodman et al., 2021, Blazev et al., 2022, Blazev et al., 2021, Molendijk et al., 2022). The precursor MS tolerance was set to 20 ppm and the MS/MS tolerance was set to 0.02 Da with a maximum of 2 miss-cleavage. The peptides were searched with oxidation of methionine set as variable modification, and carbamidomethylation of cysteine and TMT to peptide N-termini and Lysine set as a fixed modification set as a fixed modification. Peptide spectral matches were filtered to 1% FDR using a target/decoy approach with Percolator (17952086). The grouped peptide data was further filtered to 1% protein FDR using Protein Validator. Quantification was performed with the reporter ion quantification node for TMT quantification in Proteome Discoverer. TMT precision was set to 20 ppm and corrected for isotopic impurities. Only spectra with < 50% co-isolation interference were used for quantification with an average signal-to-noise filter of > 10.

XL-MS data were searched against the same skeletal muscle mouse specific proteome database with pLink2 (v2.3.11, Chen et al., 2019). The peptides were searched with oxidation of methionine set as variable modification, and carbamidomethylation of cysteine and TMT to peptide N-termini and Lysine set as a fixed modification set as a fixed modification. Default parameters were used with up to 2 missed cleavages allowed, filter tolerance was set to 10 ppm and peptides were filtered to 1% FDR. XL-MS search was performed using tBu-PhoX as the set linker, where elemental composition was manually imported using cross-linker information from ProteomeDiscoverer. Extraction of reporter ion intensities was performed using an inhouse generated script. Briefly, the msaccess tool from ProteoWizard (Chambers et al., 2012) was used to extract selected ion chromatograms for the masses shown in **Table 1** below, with a 10 ppm radius. Peak intensity data for each reporter ion were merged by scan index and peak identifier before exporting as a table for further analysis.

**Table 1.** TMT masses for extracting ion intensity.

| Mass | Label | ppm |
| --- | --- | --- |
| 126.127726 | TMTpro-126 | 10 |
| 127.124761 | TMTpro-127N | 10 |
| 127.131081 | TMTpro-127C | 10 |
| 128.128116 | TMTpro-128N | 10 |
| 128.134436 | TMTpro-128C | 10 |
| 129.131471 | TMTpro-129N | 10 |
| 129.13779 | TMTpro-129C | 10 |
| 130.134825 | TMTpro-130N | 10 |
| 130.141145 | TMTpro-130C | 10 |
| 131.13818 | TMTpro-131N | 10 |
| 131.1445 | TMTpro-131C | 10 |
| 132.141535 | TMTpro-132N | 10 |
| 132.147855 | TMTpro-132C | 10 |
| 133.14489 | TMTpro-133N | 10 |
| 133.15121 | TMTpro-133C | 10 |
| 134.148245 | TMTpro-134N | 10 |
| 134.154565 | TMTpro-134C | 10 |
| 135.1516 | TMTpro-135N | 10 |

PCP-MS DIA data were processed with Spectronaut v18.6.231127 against the mouse skeletal muscle database using DirectDIA with default parameters with precursor and protein Qvalue cutoff set to 0.01. Peptide quantification was carried out at MS2 level using 3-6 fragment ions, with automatic interference fragment ion removal as previously described (Bruderer et al., 2017). Dynamic MS1 and MS2 mass tolerance was enabled, and retention time calibration was accomplished using local (non-linear) regression. The default dynamic extracted ion chromatogram window size was performed.

IodoTMT data was searched against the mouse skeletal muscle database with SequestHT within Proteome Discoverer v2.5.0.4 (Eng et al., 1994). The precursor MS tolerance was set to 20 ppm and the MS/MS tolerance was set to 0.02 Da with a maximum of 2 miss-cleavage. The peptides were searched with oxidation of methionine, iodoTMT and Nethylmaleimide of Cysteine set as variable modification. Peptide spectral matches were filtered to 1% FDR using a target/decoy approach with Percolator (Kall et al., 2007). Quantification was performed with the reporter ion quantification node for TMT quantification in Proteome Discoverer. TMT precision was set to 20 ppm and corrected for isotopic impurities. Only spectra with < 50% co-isolation interference were used for quantification with an average signal-to-noise filter of > 10.

### Complexomic analysis using CCProfiler

Peptide level data was imported into the CCProfiler Analysis Package (Bludau et al., 2023). Non-proteotypic peptides were filtered out and molecular weight calibration of fractions was annotated. Protein expression was then converted to a traces list format where each peptide trace was annotated with Uniprot protein ID, entry name and molecular weight. Missing peptide intensity values (for which both the previous and the following fraction contained measured intensity values) were imputed by a spline fit across the fractions. After missing value imputation, peptide intensity values were normalized across conditions and replicates by applying a cyclic loess normalization (Rosenberger et al., 2020, Bolstad et al., 2003, Ballman et al., 2004) a method initially implemented for microarray analysis using pairwise loess curve fitting cycling through all possible pairs several times. Low-confidence peptides were subsequently removed, keeping only peptides with (1) at least three consecutive detections across any replicate, (2) at least one high correlating sibling peptide (maximum correlation ≥ 0.5), and (3) a good average sibling peptide correlation (≥0.2). Protein quantification was performed by summing the top two most intense peptides consistently across all replicates. Protein level data was then quantified using the peptide traces and subject to protein centric and complex centric modules of differential analysis. Protein-centric analysis was performed with following parameters: corr_cutoff = 0.7, window_size = 7, rt_height = 1, smoothing_length = 7, perturb_cutoff = “5%”, and collapse_method = “apex_only”.

For complex-centric analysis, we first defined a set of target protein complex queries. This was achieved by combining queries derived from CORUM (Ruepp et al., 2010, Tsitsiridis et al., 2023), StringDB (Snel et al., 2000) and hu.MAP.3 (Fischer et al., 2024). We derived protein complex queries from StringDB version 10 (10090.protein.links.v12.txt). Protein identifiers were mapped to Uniprot accessions via BioMart. The interactions were filtered for a minimal combined_score of 980. We applied the CCProfiler clustering algorithm to generate stringDB based complex queries. Mouse CORUM and human hu.MAP.3 derived protein complex queries were taken, the complex queries were combined, and decoys were generated randomly by requiring a minimum edge distance of 3. Complex-centric analysis was performed with the following parameters: corr_cutoff = 0.9, window_size = 7, rt_height = 1, smoothing_length = 7, perturb_cutoff = “5%” and collapse_method = “apex_network”. Only complex features with a molecular weight higher than two times the largest monomeric molecular weight of any of its participating subunits were considered. For each protein complex query, the complex feature with the highest number of participating subunits was selected for the FDR estimation, filtering for a maximum FDR of 5%. Secondary features were appended to the final results based on the basis of a minimum peak correlation threshold of 0.7. To reduce redundancy across the detected complex features between different queries, features were collapsed with the following parameters: rt_height = 0 and distance_cutoff = 1.25.

### Pairwise protein correlation analysis

A pairwise correlation analysis was performed on normalized protein intensities across the BN-PAGE fractions where a Pearson correlation coefficient was generated for all protein pairs.

### Quantification & Statistical Analysis

For global total proteomics, XL-MS and cysteine-redox proteomics, log_2_(x) transformation and median subtraction normalization was performed in Perseus (Tyanova et al., 2016). Differential abundance was calculated with Student’s t-test and q-values generated using Benjamini-Hochberg with FDR set to 5%. Pathway enrichment analysis was performed with TeaProt (Molendijk et al., 2023).

### Data availability

Raw data is available for download from the University of Melbourne Mediaflux:

**PCP-MS analysis of insulin resistance in C2C12 myotubes with or without acute insulin stimulation:** https://mediaflux.researchsoftware.unimelb.edu.au/mflux/data/mover/index.html?token=kg48rxcqq76v15kvwwsfj2iqkvmijugyydjrqr2jq8gseyh4ojrt1nw0rg3dq2k70l23lv5klxose7z8nrm7eccvx47plvyt340yrptojjxjssiixva7joospoujpxh3lc4bf3uma617ngv067hnz2b8oxosub91p29j4mgnc3cakxv4xg5ojoienj1tnm58hmkr7egzbiqjt7zqlamt3wmnl3p7rkfz4pp9mt7y

**qXL-MS analysis of insulin resistance in C2C12 myotubes with or without acute insulin stimulation:** https://mediaflux.researchsoftware.unimelb.edu.au/mflux/data/mover/index.html?token=kg48rxcqq76v15kvwwsfj2iqkvmijugyydjrqr2jq8gseyh4ojrt1nw0rg3dq2k70l23lv5klxose7z8nrm7eccvx47plvyt2l42pz49rcgzme5u2ousxkdwmmgvjaaxidtxyfxq7acozilhp4gpv4ctf18p28icydmo8aq0oj6kp26ggptm7eywcmuf7y0qtz30mjuqo58fwihb3acj33z1c337y7n9j94dk5yr

**qXL-MS analysis of insulin resistance in mouse skeletal muscle:** https://mediaflux.researchsoftware.unimelb.edu.au/mflux/data/mover/index.html?token=kg48rxcqq76v15kvwwsfj2iqkvmijugyydjrqr2jq8gseyh4ojrt1nw0rg3dq2k70l23lv5klxose7z8nrm7eccvx47plvyr8nfmkycrp3zvb2nd3135v0behl1o9vzbb83k5lud4lv3fd29sp767etlr4kwqzw908fi48dusblvywg6m9ujf76eghltls3anpbbvjoqwbzgnb7eq5a0i7hsn7s0hh6ac3962d0r

**Cysteine Redox proteomics of PDIA6 over-expressing skeletal muscle:** https://mediaflux.researchsoftware.unimelb.edu.au/mflux/data/mover/index.html?token=kg48rxcqq76v15kvwwsfj2iqkvmijugyydjrqr2jq8gseyh4ojrt1nw0rg3dq2k70l23lv5klxose7z8nrm7eccvx47plvynkro576l4xkbfmw24v2p5crdkcwkiwy7q31p6mwgkou99xuciyh7fht5248q0bae5jaf3o8htj1llnjnuvlsg0qb8abyiz5c615smk9sg0vsp5wz7i97dun9onk49ipb63b5c9u2f

## Notes

https://mediaflux.researchsoftware.unimelb.edu.au/mflux/data/mover/index.html?token=kg48rxcqq76v15kvwwsfj2iqkvmijugyydjrqr2jq8gseyh4ojrt1nw0rg3dq2k70l23lv5klxose7z8nrm7eccvx47plvyt340yrptojjxjssiixva7joospoujpxh3lc4bf3uma617ngv067hnz2b8oxosub91p29j4mgnc3cakxv4xg5ojoienj1tnm58hmkr7egzbiqjt7zqlamt3wmnl3p7rkfz4pp9mt7y

https://mediaflux.researchsoftware.unimelb.edu.au/mflux/data/mover/index.html?token=kg48rxcqq76v15kvwwsfj2iqkvmijugyydjrqr2jq8gseyh4ojrt1nw0rg3dq2k70l23lv5klxose7z8nrm7eccvx47plvyt2l42pz49rcgzme5u2ousxkdwmmgvjaaxidtxyfxq7acozilhp4gpv4ctf18p28icydmo8aq0oj6kp26ggptm7eywcmuf7y0qtz30mjuqo58fwihb3acj33z1c337y7n9j94dk5yr

https://mediaflux.researchsoftware.unimelb.edu.au/mflux/data/mover/index.html?token=kg48rxcqq76v15kvwwsfj2iqkvmijugyydjrqr2jq8gseyh4ojrt1nw0rg3dq2k70l23lv5klxose7z8nrm7eccvx47plvyr8nfmkycrp3zvb2nd3135v0behl1o9vzbb83k5lud4lv3fd29sp767etlr4kwqzw908fi48dusblvywg6m9ujf76eghltls3anpbbvjoqwbzgnb7eq5a0i7hsn7s0hh6ac3962d0r

https://mediaflux.researchsoftware.unimelb.edu.au/mflux/data/mover/index.html?token=kg48rxcqq76v15kvwwsfj2iqkvmijugyydjrqr2jq8gseyh4ojrt1nw0rg3dq2k70l23lv5klxose7z8nrm7eccvx47plvynkro576l4xkbfmw24v2p5crdkcwkiwy7q31p6mwgkou99xuciyh7fht5248q0bae5jaf3o8htj1llnjnuvlsg0qb8abyiz5c615smk9sg0vsp5wz7i97dun9onk49ipb63b5c9u2f

### Competing Interest Statement

The authors have declared no competing interest.

